# Grapevine pruning time affects natural wound colonization by wood-invading fungi

**DOI:** 10.1101/2020.04.20.050955

**Authors:** María del Pilar Martínez-Diz, Ales Eichmeier, Milan Spetik, Rebeca Bujanda, Ángela Díaz-Fernández, Emilia Díaz-Losada, David Gramaje

**Author notes:** Corresponding author. Instituto de Ciencias de la Vid y del Vino (ICVV), Consejo Superior de Investigaciones Científicas, Universidad de la Rioja, Gobierno de La Rioja, Ctra. LO-20 Salida 13, 26007 Logroño, Spain. E-mail address (D. Gramaje).

## Abstract

Grapevine pruning wounds made during the dormant season are a port of entry of wood-invading fungi. Timing of pruning may affect the wound susceptibility to these fungi, such as those associated with grapevine trunk diseases (GTDs). This study aimed to determine the effect of pruning time on natural fungal infection in six vineyards in Galicia, Spain, belonging to three Denominations of Origin (D.O) over two growing seasons. Pruning wounds were left unprotected physically and chemically during two periods of three months each, from November to February and from February to May. The diversity and composition of the fungal microbiome that colonized the pruning wounds was identified by ITS2 high-throughput amplicon sequencing (HTAS). A broad range of fungi was able to colonize grapevine pruning wounds at both infection periods. Fungal microbiome composition did not shift as year of sampling. Fungal communities were affected in their diversity and composition by the D.O., whereas the spatial variation (i.e. vineyard within each region) was low. Pruned canes harboured a core community of fungal species, which appeared to be independent of the infection period. Accumulated rainfall over 8 and 11 weeks after pruning positively correlated with the total fungal microbiome and in particular with the GTD fungal genus *Diaporthe* abundances. A strong seasonal effect on GTD fungal infection was detected for most genera, with higher percentages of abundance detected after pruning in February (winter) as compared with that of pruning in November (mid-autumn). In light of the GTD colonization results and given the environmental conditions and the geographical location of this study, early pruning is recommended to reduce the infections caused by GTD fungi during the pruning season in Galicia.

## 1. Introduction

The most important operation during the dormant season in vineyards is pruning. Pruning of grapevines is recommended any time after leaf fall, which may occur late fall or throughout the winter. The purpose of pruning is to obtain maximum yields of high-quality grapes and to allow adequate vegetative growth for the following season (Jackson, 2004). Timing of pruning within the dormant season may affect the grapevine phenology, and thus yield and fruit quality (Zheng et al., 2017). Early or late pruning can also affect the susceptibility of the plant to abiotic disorders, such as spring frost (Jackson, 2004), or the pruning wound susceptibility to infections caused by wood-invading fungi, such as those associated with grapevine trunk diseases (GTDs) (Luque et al., 2014).

Grapevine trunk diseases are caused by a broad range of taxonomically unrelated fungal pathogens that infect woody tissues. They reduce longevity and productivity of grapevines and thereby cause substantial economic losses to industry (Gramaje et al., 2018). To date, up to 135 fungal species belonging to 35 genera have been associated with GTDs worldwide (Gramaje et al., 2018; Aigon-Mouhous et al., 2019; Lawrence et al., 2019; Berlanas et al., 2020), thus accounting for the largest group of fungi known to infect grapevines (Gramaje et al., 2018). GTDs are mainly caused by fungal ascomycetes but some basideomiceteous fungi are also thought to play a relevant role in this pathosystem (Fischer, 2002; Cloete et al., 2015; Brown et al., 2020). GTD fungal spores can infect grapevines through any open wound, including those caused by de-suckering, trimming and re-training (Makatini et al., 2014). Nonetheless, annual pruning wounds are the primary point of infection providing many entry sites each growing season throughout the life of a vineyard (Gramaje et al., 2018).

The main GTDs in mature vines are Eutypa dieback, Botryosphaeria dieback, Phomopsis dieback and esca disease (Gramaje et al., 2018). In North America, several *Cytospora* spp. have also been recently reported causing dieback and wood cankers in grapevine (Lawrence et al., 2017). Grapevine pathogens responsible for these diseases are mainly spread through the dispersion of airborne spores. Previous studies showed that spore release and thus, high risk periods of infection vary during the growing season depending on the geographical location and fungal pathogen but mainly overlay with dormant pruning seasons in both the Southern and Northern Hemispheres (Pearson, 1980; Petzoldt et al., 1983; Eskalen and Gubler, 2001; Amposah et al., 2009; Trouillas, 2009; Úrbez-Torres et al., 2010; van Niekerk et al., 2010; Valencia et al., 2015). Pruning wounds susceptibility to GTD pathogens primarily depends on the pruning time and the period elapsed between pruning and possible infection cases. Studies using artificial spore inoculations indicate that susceptibility of grapevine pruning wound significantly decreases as the length of time between pruning and inoculation increases, with seasonal variation noted between regions, due mainly to climatic differences (Moller and Kasimatis, 1980; Munkvold and Marois, 1995; Eskalen et al., 2007; Serra et al., 2008; Úrbez-Torres and Gubler, 2011; van Niekerk et al., 2011; Ayres et al., 2016).

The rate of natural fungal microbiome infections in pruned canes has been poorly studied so far, and data available is only referred in the context of GTD pathogens infections in France (Lecomte and Bailey, 2011) and northeast Spain (Luque et al., 2014). These studies employed standard culture-dependent microbial techniques; however, these approaches tend to underestimate species richness and misrepresent fungal activity, because fungi may be highly selective, hidden and slow growing. Molecular-based methods have progressively replaced morphological techniques to characterize the microbiome in nature. These methods allow the detection and identification of a greater number of microorganisms, including species that are unable to be isolated in culture (Amann et al., 1995). The novel advances in high-throughput sequencing (HTS) technology have increased both the resolution and scope of fungal community analyses and have revealed a highly diverse and complex fungal microbiome of plant vascular systems (Studholme et al., 2011).

In recent years, grapevine has become a plant model system for microbiome research. HTS tools have been actively used to map the microbiome on grapevine organ epiphytes (i.e., root, leaf and berry) because of its importance with grape production and specially with regards to foliar and fruit diseases control along with the biological implication of indigenous microorganisms with the local signature of a wine (Bokulich et al., 2014; Perazzolli et al., 2014; Zarraonaindia et al., 2015). Identification of the microbial communities inhabiting the grapevine endosphere has been achieved using standard culture-dependent microbial techniques (West et al., 2010; Compant et al., 2011; Baldan et al., 2014; Kraus et al., 2019). Culture-independent high-throughput amplicon sequencing (HTAS) approaches have recently been used to improve the microbiome profiling of grapevine woody organs such as cane and trunk (Faist et al., 2016; Deyett et al., 2017; Dissanayake et al., 2018; Eichmeier et al., 2018).

In this study, we tested the following hypotheses: (1) the diversity and composition of fungal microbiome that colonizes grapevine pruning wounds changes according to the pruning time and this shift is related to environmental conditions; (2) the susceptibility of pruning wounds to fungal infection and the ability of GTD pathogens to colonize them depend on the pruning time, therefore this would allow us to make pruning recommendations to growers in the short term in order to avoid high pathogen infection periods. The objective was therefore to identify the diversity and composition of the fungal microbiome, in particular GTD fungi, infecting pruning wounds in six mature vineyards at two pruning times over two years by HTAS: (i) after an early pruning in autumn; and (ii) after a late pruning in winter. In addition, we investigated the relationship between the main weather data recorded during the experimental period and the rate of fungal colonization.

## 2. Materials and methods

### 2.1 Location and characteristics of the experimental vineyards

Experiments were conducted at six experimental plots located in three Denominations of Origin (D.O. Valdeorras, D.O. Ribeiro and D.O. Rías Baixas; two experimental plots per D.O.) in Galicia region, Spain (Table S1), from November 2017 to May 2019. Plots within each D.O. were <10 km apart and had very similar climates. Standard cultural practices were employed in all sites during the grapevine growing season, and the management of powdery and downy mildews was performed using only wettable sulphur and copper compounds and applied at label dosages and following IPM guidelines, respectively, if required. Plots of 1,500 vines in these vineyards have been monitored biannually for the evolution of GTD symptoms since 2014 to the present. At the time this study was started (2017), about 12% of vines had shown symptoms of trunk diseases in previous monitoring dates. The main symptoms of GTDs observed during monitoring included chlorotic leaves, stunted shoots, and short internodes (Eutypa dieback), the arm and cordon death (Botryosphaeria, Eutypa and Phomopsis diebacks) and tiger-pattern foliar necrosis (esca). All vineyards were trained as bilateral cordons with spur pruning (Royat). Grapevine cultivars differed among D.O. (Table S1), so data from each D.O. was analysed independently due to the previously reported variable degree of susceptibility of each grapevine cultivar to fungal trunk pathogen infections (Martínez-Diz et al., 2019a). The experimental plots were located <6km to automatic weather stations owned by MeteoGalicia (Weather Service of Galician Regional Government, Xunta de Galicia). Data obtained from the weather station in each D.O. were considered to be representative of the two experimental plots.

### 2.2 Pruning and sampling

A total of 200 vines were pruned in each experimental plot in mid-autumn (between 13 and 14 November for both years and experimental plots) leaving six buds. Then, 25 pruned canes in each vineyard were chosen at random and labelled for subsequent samplings. Wood of these 25 canes were taken to the laboratory for DNA extraction. Three months later in winter (between 21 and 22 February for both years and experimental plots), a 15-cm section was cut from the labelled 25 pruned canes and removed from their upper end and taken to the laboratory for DNA extraction. On the same day of this sampling, all the vines were pruned to four buds. A longer than usual wood section (5–7 cm) was left above the top bud. Three months later in spring (between 22 and 23 May for both years and experimental plots), sampling for DNA extraction from approximately a 15-cm wood section was repeated following the same procedure earlier, and the labelled canes were definitely pruned to two buds. All canes were therefore exposed to natural infections for three months after pruning (infection period 1: November-February; infection period 2: February-May). Pruning scissors were disinfested with 70% ethanol every pruning cut. Pruning wounds were not protected physically nor chemically during the experiment.

### 2.3 DNA extraction and sequencing

Before DNA extraction, pruned canes were sequentially washed in 70% ethanol and sterile distilled water. Upon this treatment, bark was carefully peeled out from the upper ends of canes with a flame-sterilised scalpel to expose the inner tissues starting from the pruning wound. The 3-mm end was cut and discarded to avoid bias by the colonization of saprophytic fungal species. DNA was extracted from 0.5 g of xylem tissue collected between 3- to 8-mm from the pruning wound using the i-genomic Plant DNA Extraction Mini Kit (Intron Biotechnology, South Korea). DNA yields from each sample were quantified using the Invitrogen Qubit 4 Fluorometer with Qubit dsDNA HS Assay (Thermo Fisher Scientific, Waltham, USA), and the extracts were adjusted to 10-15 ng/µl. After DNA quantification, samples of each pruning time and vineyard were pooled in groups of five, resulting in a total of five replicates for every batch of 25 canes. A total of 180 DNA samples was analysed. Complete fungal ITS2 region (around 300 bp) was amplified using the primers ITS86F (5’ GTGAAT CATCGAATCTTTGAA 3’) (Turenne et al., 1999) and ITS4 (5’ TCCTCCGCTTATTGATATGC 3’) (White et al., 1990), to which the Illumina sequencing primer sequences were attached to their 5’ ends. PCRs were carried out in a final volume of 25 µL, containing 2.5 µL of template DNA, 0.5 µM of the primers, 12.5 µL of Supreme NZYTaq 2x Green Master Mix (NZYTech, Lisboa, Portugal), and ultrapure water up to 25 µL. The reaction mixture was incubated as follows: initial denaturation at 95 oC for 5 min, followed by 35 cycles of 95 oC for 30 s, 49 oC for 30 s, 72 oC for 30 s, and a final extension step at 72 oC for 10 minutes. The oligonucleotide indices which are required for multiplexing different libraries in the same sequencing pool were attached in a second PCR round with identical conditions but only five cycles and 60 oC as the annealing temperature for a schematic overview of the library preparation process. A negative control that contained no DNA was included in every PCR round to check for contamination during library preparation (BPCR). The libraries were run on 2 % agarose gels stained with GreenSafe (NZYTech, Lisboa, Portugal) and imaged under UV light to verify the library size. Libraries were purified using the Mag-Bind RXNPure Plus magnetic beads (Omega Biotek, Norcross, GA, USA), following the instructions provided by the manufacturer. Then, they were pooled in equimolar amounts according to the quantification data provided by the Qubit dsDNA HS Assay (Thermo Fisher Scientific, Waltham, USA). The pool was sequenced in a MiSeq PE300 run (Illumina, San Diego, USA). An additional negative control was included during the extraction step. A positive control containing DNA of a grapevine endorhizosphere sample previously evaluated by ITS HTAS was also included (Martínez-Diz et al., 2019b). All control samples were prepared for sequencing to evaluate potential contaminations of the entire process.

### 2.4 Data analysis of the high-throughput amplification assay

Sequence quality was visualized using FastQC-0.10.1 (Andrews, 2010), and the CLC Genomics Workbench 6.5.1 (CLC Bio, Aarhus, Denmark) was used to trim and merge the paired end reads. The parameter Q30 was applied and only reads longer than 100 nts with average read quality > 30 were considered for further analysis. Q30 represents the quality score of a base, also known as a Phred or Q score. It is an integer value representing the estimated probability of an error. Q30 number means base call accuracy 99.9%. The distance of evaluated reads in the trimming and merging step was set from 200 to 400 nts. Primer and Illumina adapter sequences were also trimmed out. The reads were exported to fasta format by CLC Genomics Workbench 6.5.1 (CLC Bio, Aarhus, Denmark).

Exported fasta files were used for clustering in SCATA (https://scata.mykopat.slu.se/). Parameters for clustering were: Clustering distance 0.015; Minimum alignment to consider clustering 0.95; Missmatch penalty 0.1; Gap open penalty 0; Gap extension penalty 1; End gap weight 0; Collapse homopolymers 3; Downsample sample size 0; Remove low frequency genotypes 0; Tag-by-Cluster Max 10000000; Blast E-value cutoff 1e-60; Cluster engine USERACH; Number of repseqs to report 50. The CBS isolates were used as a reference sequences. Singleton operational taxonomic units (OTUs) were discarded. The sequences of non-singleton OTUs were used as the representative sequence and were identified using the blastn algorithm using the GenBank/NCBI reference database (version 2.2.30+). OTUs with no kingdom-level classification or matching chloroplast, mitochondrial or Viridiplantae sequences were excluded from the dataset. In order to optimize the dataset, each sample was rarefied to the same sequence number per sample, that is, 21,287 fungal sequences. OTUs represented globally by less than five reads were discarded (Glynou et al., 2018). The resulting quality dataset was used for the estimation of richness and diversity. The metadata, OTUs table and associated taxonomic classifications deployed in this study have been deposited in figshare (ID: 79113). HTAS data were deposited in GenBank/NCBI under BioProject Acc. No. PRJNA625395.

### 2.5 Fungal diversity, taxonomy distribution and statistical analysis

Within sample type, alpha-diversity estimates were calculated by analyzing the Shannon diversity and Chao1 richness in Phyloseq package, as realized in the tool MicrobiomeAnalyst (Dhariwal et al., 2017). Multiple mean comparisons using Tukey’s test were performed to determine how fungal alpha-diversity differed among year, D.O., vineyard within each D.O., and pruning time. *P* values were corrected for multiple comparisons using the sequential Bonferroni correction. Relationship in OTUs composition among samples were investigated by calculating Bray Curtis metrics, and visualized by means of PCoA plots (Vázquez-Baeza et al., 2013) using MicrobiomeAnalyst. PERMANOVA was performed to investigate which OTUs significantly differed in abundance among experimental factors. Rarefaction curves and Good’s coverage values were calculated using MicrobiomeAnalyst.

The Linear Discriminant Analysis Effect Size (LEfSe) algorithm was used to identify taxa (genus level or higher) that differed in relative abundance between pruning times (Segata et al., 2011). The online MicrobiomeAnalyst interface was used, the threshold for the logarithmic Linear Discriminant Analysis (LDA) score was set at 1.0 and the Wilcoxon *p*-value at 0.05. The results are displayed in a bar graph. The fungal OTUs shared among compartments were obtained by a Venn-diagram analysis using the software retrieved from http://bioinformatics.psb.ugent.be.

In order to compare the percentage of abundance of each fungal genus associated with GTDs between both infection periods, an analysis of variance with log transforms was used. Normality of residuals was checked by Shapiro-Wilk’s test, and homogeneity of variances by Levene’s test. Means were compared with Tukey’s Honestly Significant Difference range test (*P*≤ 0.05) using Statistix 10 software (Analytical Software).

### 2.6 Correlation with weather variables

The number of OTUs corresponding to the total fungal microbiome, and the fungal genera associated with GTDs was correlated with the main weather data (daily mean relative humidity, daily mean temperature and accumulated rainfall). Values from the number of OTUs were transformed by log (n/N * 1000 + 1). Where n was the number of OTUs detected on each sample and N was the total number of OTUs detected. Temperature and humidity records were averaged over 1, 2, 4, 8, and 11 weeks post-pruning periods. Rainfall records were accumulated and log-transformed to make data conform to normality over the same periods. Spearman’s correlation coefficients were calculated using the function *cor* of the ‘stats’ package of R v. 3.6.0 (R Core Team, 2019).

## 3. Results

### 3.1 High-throughput amplicon sequencing

After paired-end alignments, quality filtering and deletion of chimeras, singletons, a total of 10,740,761 fungal internal transcribed spacer (ITS2) sequences were generated from 180 samples, excluding controls, and assigned to 259 fungal OTUs.

OTUs generated from the negative control used in the amplification step belonged to 30 genera. The number of sequences of each OTU present in the negative control was subtracted from the sequence abundance of that OTU in the experimental samples according to Nguyen et al. (2015). No contamination was detected in the negative control used in the DNA extraction step. Good’s coverage values in all samples ranged from 99.25 to 100%, capturing nearly all the diversity with an adequate sequencing depth (Figure S1). Chao1 diversity estimator ranged from 5 to 26, while Shannon diversity estimator ranged from 0.31 to 2.27 (Table S2).

### 3.2 Fungal communities differed among Denominations of Origin

The alpha-diversity of fungal communities differed among D.O. (Chao1: *P* = 0.0047, Shannon: *P* < 0.001; Fig 1), and principal coordinates analysis (PCoA) of Bray Curtis data demonstrated that D.O. was the primary source of beta-diversity (*R*^2^ = 0.48, *P* < 0.001) (Fig. 2). Therefore, data of each D.O. was analysed independently.

**Figure 1.**
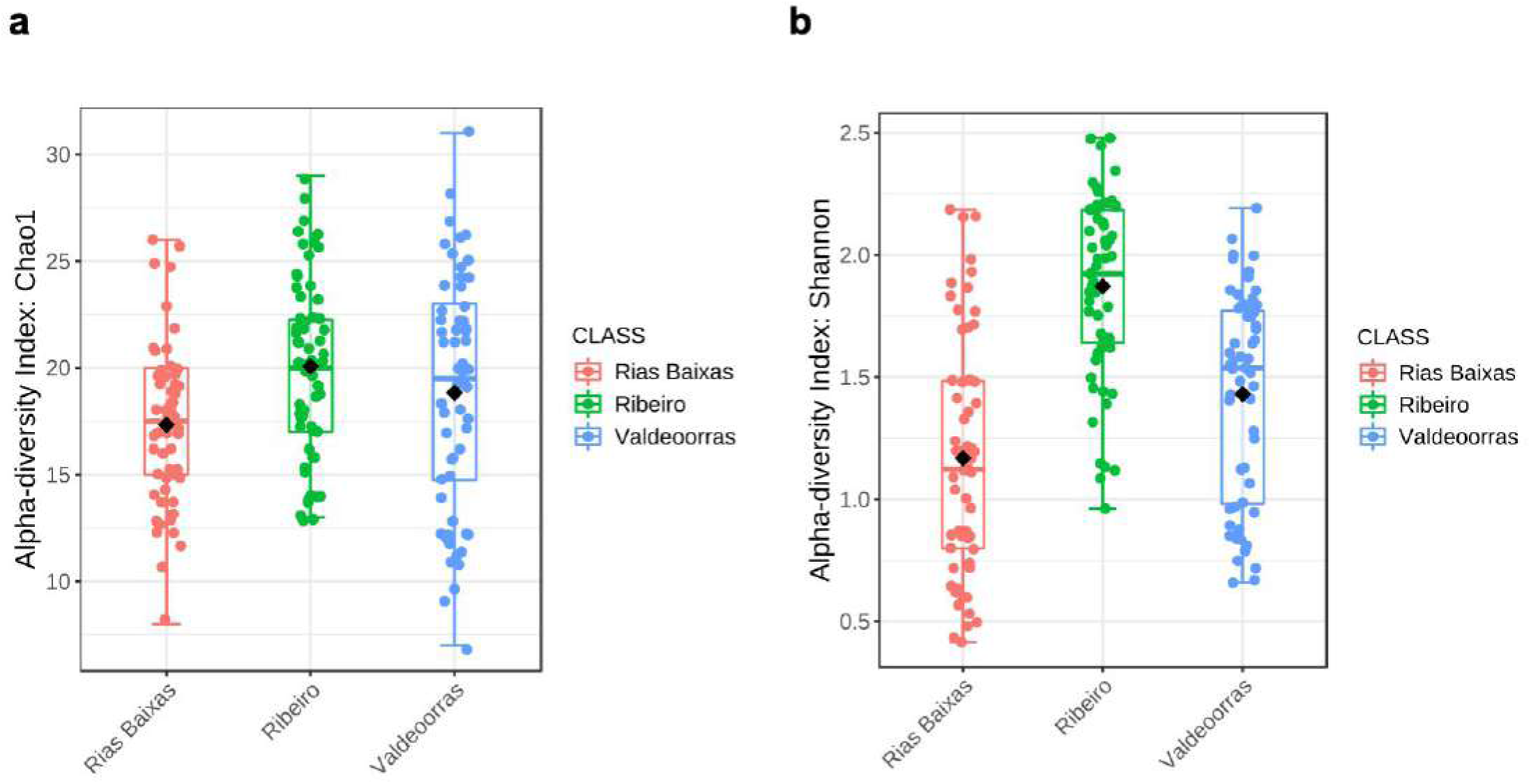
Boxplot illustrating the differences in Chao1 **(a)** and Shannon **(b)** diversity measures of the fungal communities in the three Denominations of Origin.

**Figure 2.**
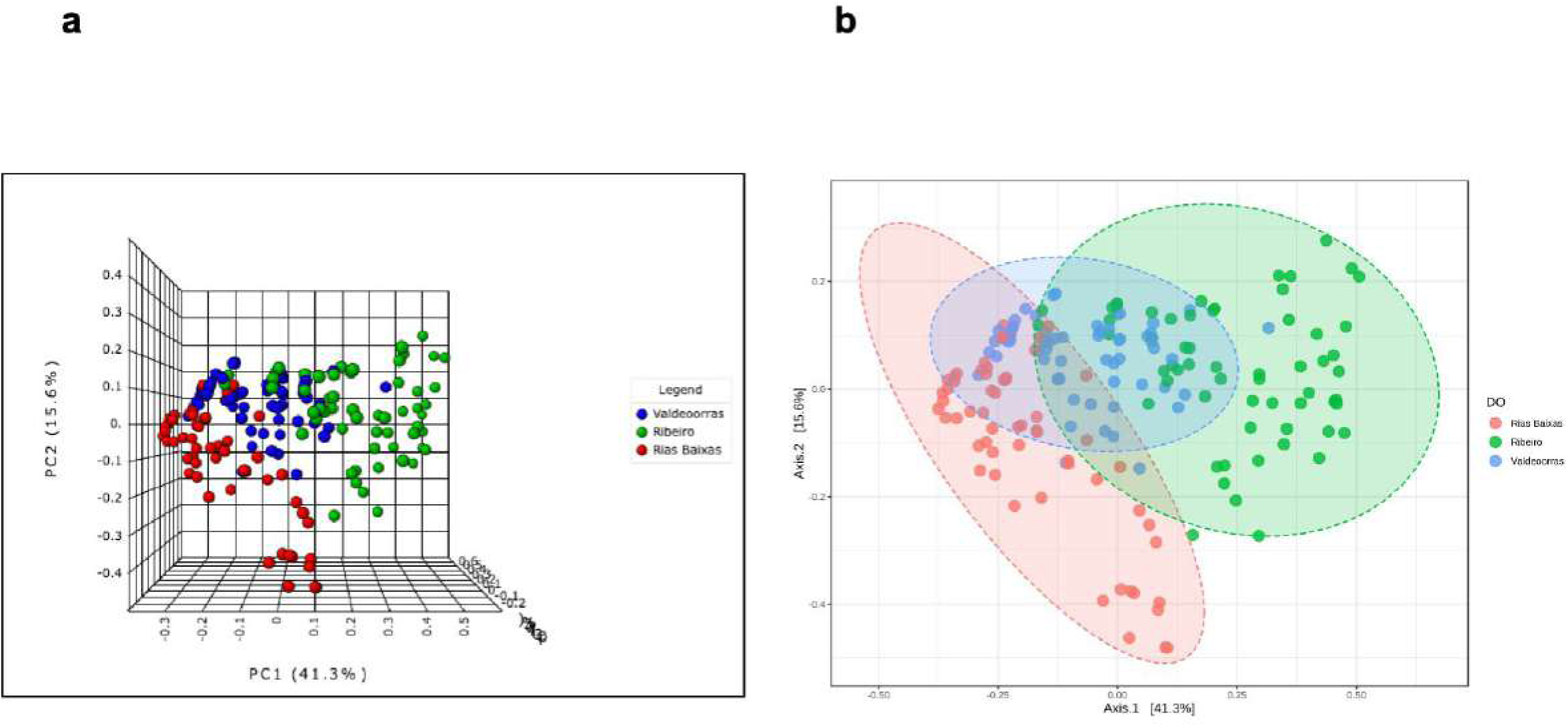
Principal Coordinate Analysis (PCoA) based on Bray Curtis dissimilarity metrics in 3D **(a)** and 2D **(b)**, showing the distance in the fungal communities among Denominations of Origin.

The relative abundance of fungal phyla, order and family detected across all D.O. is shown in Fig. S2. Considering the three D.O., the most abundant phyla were Ascomycota, followed by Basidiomycota (Fig. S2a). The most abundant orders were Dothideales, followed by Capnodiales and Pleosporales (Fig. S2b). The most abundant families were Dothioraceae, followed by Cladosporiaceae and Dermateaceae (Fig. S2c). Comparing the fungal microbiota of the three D.O., 56.8% of fungal OTUs were shared among them (Fig. 3). Specific OTUs associated with each vineyard ranged from 12.1 to 18.4% of their fungal communities (Fig. 3).

**Figure 3.**
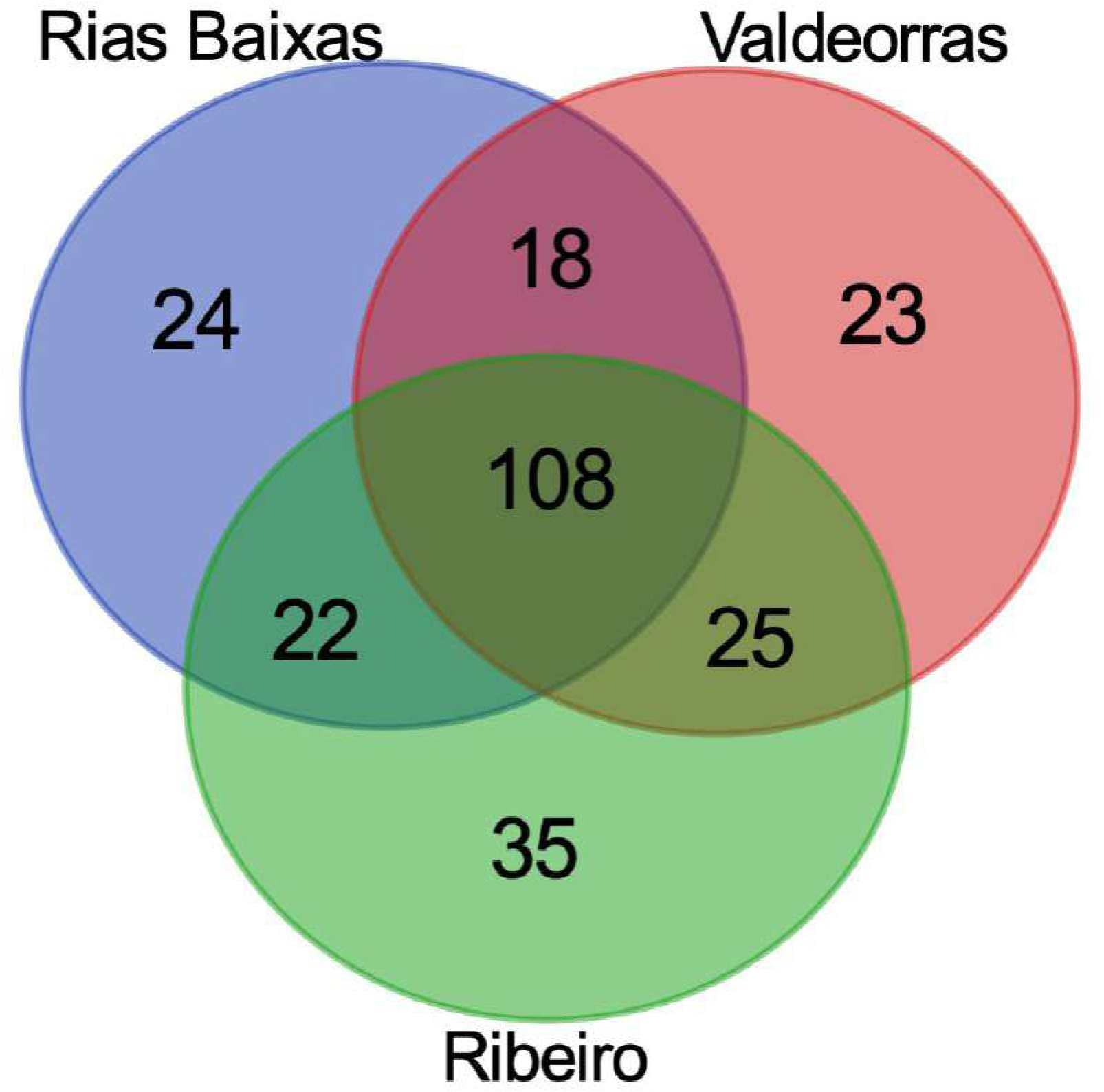
Venn diagram illustrating the overlap of the OTUs identified in the fungal microbiota among Denominations of Origin.

In D.O. Rías Baixas, the most abundant families were Dothioraceae (62.9%), followed by Cladosporiaceae (10.2%) and Pleosporaceae (9.9%) (Fig. S2c). The most abundant families in D.O. Ribeiro were Cladosporiaceae (26.5%), followed by Dothioraceae (21.6%) and Dermateaceae (12.1%). In D.O. Valdeorras, the most abundant families were Dothioraceae (51.8%), followed by Cladosporiaceae (19.9%) and Tremellaceae (4.5%).

### 3.3 Fungal diversity exhibits a temporal variation over the infection periods

Alpha-diversity of fungal communities in grapevine wood samples did not differ significantly between experimental plots (Fig. S3) and year (Fig. S4) within each D.O. (Table 1), thus the data of both years and experimental plots were combined for analyses. Comparing the microbiome in the grapevine inner tissue at the three sampling times (1: November, 2: February and 3: May), higher fungal diversity was mostly observed towards the sampling time 3 (*P* < 0.05) (Fig. S5). Excluding the initial fungal microbiome estimated in November, and considering the two infection periods, fungal community diversity was significantly different between both periods in D.O. Rías Baixas (Table 1; Fig. 4b). A PCoA further demonstrated that variation in the D.O. Rías Baixas dataset could be attributed to infection periods (*R*^2^ = 0.60; Fig. S6b). In D.O. Ribeiro, the infection periods did not predict Shannon diversity (Table 1; Fig. 4a), and any summary metrics of alpha-diversities in D.O. Valdeorras (Table 1; Fig. 4c). Infection periods did not affect the Bray Curtis metric of beta-diversity in D.O. Ribeiro and D.O. Valdeorras (*R*^2^ < 0.40; Fig. S6b and S6c).

**Table 1:** Experimental factors predicting alpha-diversity of pruning wounds associated fungal communities in three Denomination of Origin (D.O.) in Galicia.

|  | D.O. Ribeiro |  | D.O. Rías Baixas |  | D.O. Valdeorras |  |
| --- | --- | --- | --- | --- | --- | --- |
|  | Shannon | Chao1 | Shannon | Chao1 | Shannon | Chao1 |
| Year | $F = 3.01$<br>$P = 0.9129$ | $F = 2.99$<br>$P = 0.6998$ | $F = 2.99$<br>$P = 0.4553$ | $F = 2.79$<br>$P = 0.211$ | $F = 4.55$<br>$P = 0.1552$ | $F = 4.22$<br>$P = 0.4761$ |
| Experimental plot | $F = 2.13$<br>$P = 0.1455$ | $F = 1.26$<br>$P = 0.0647$ | $F = 3.05$<br>$P = 0.3550$ | $F = 3.77$<br>$P = 0.1295$ | $F = 3.12$<br>$P = 0.6772$ | $F = 3.79$<br>$P = 0.8890$ |
| Year x experimental plot | $F = 0.85$<br>$P = 0.8773$ | $F = 0.95$<br>$P = 0.4103$ | $F = 1.14$<br>$P = 0.1445$ | $F = 1.92$<br>$P = 0.2301$ | $F = 2.99$<br>$P = 0.9778$ | $F = 3.76$<br>$P = 0.9110$ |
| Infection period* | $F = 4.21$<br>$P = 0.7468$ | $F = 3.72$<br>**$P = 0.0416$** | $F = 3.75$<br>**$P < 0.001$** | $F = 5.06$<br>**$P < 0.001$** | $F = 4.25$<br>$P = 0.9079$ | $F = 4.20$<br>$P = 0.1221$ |
| Year x infection period | $F = 1.83$<br>$P = 0.2340$ | $F = 0.90$<br>$P = 0.0981$ | $F = 2.19$<br>$P = 0.1221$ | $F = 4.26$<br>$P = 0.2543$ | $F = 2.35$<br>$P = 0.4551$ | $F = 1.88$<br>$P = 0.2989$ |
| Experimental plot x infection period | $F = 3.24$<br>$P = 0.1134$ | $F = 4.01$<br>$P = 0.0944$ | $F = 3.11$<br>$P = 0.0987$ | $F = 5.79$<br>$P = 0.1556$ | $F = 5.67$<br>$P = 0.2375$ | $F = 4.15$<br>$P = 0.6780$ |

\*Infection period: 1 (from November to February) and 2 (from February to May)

**Figure 4.**
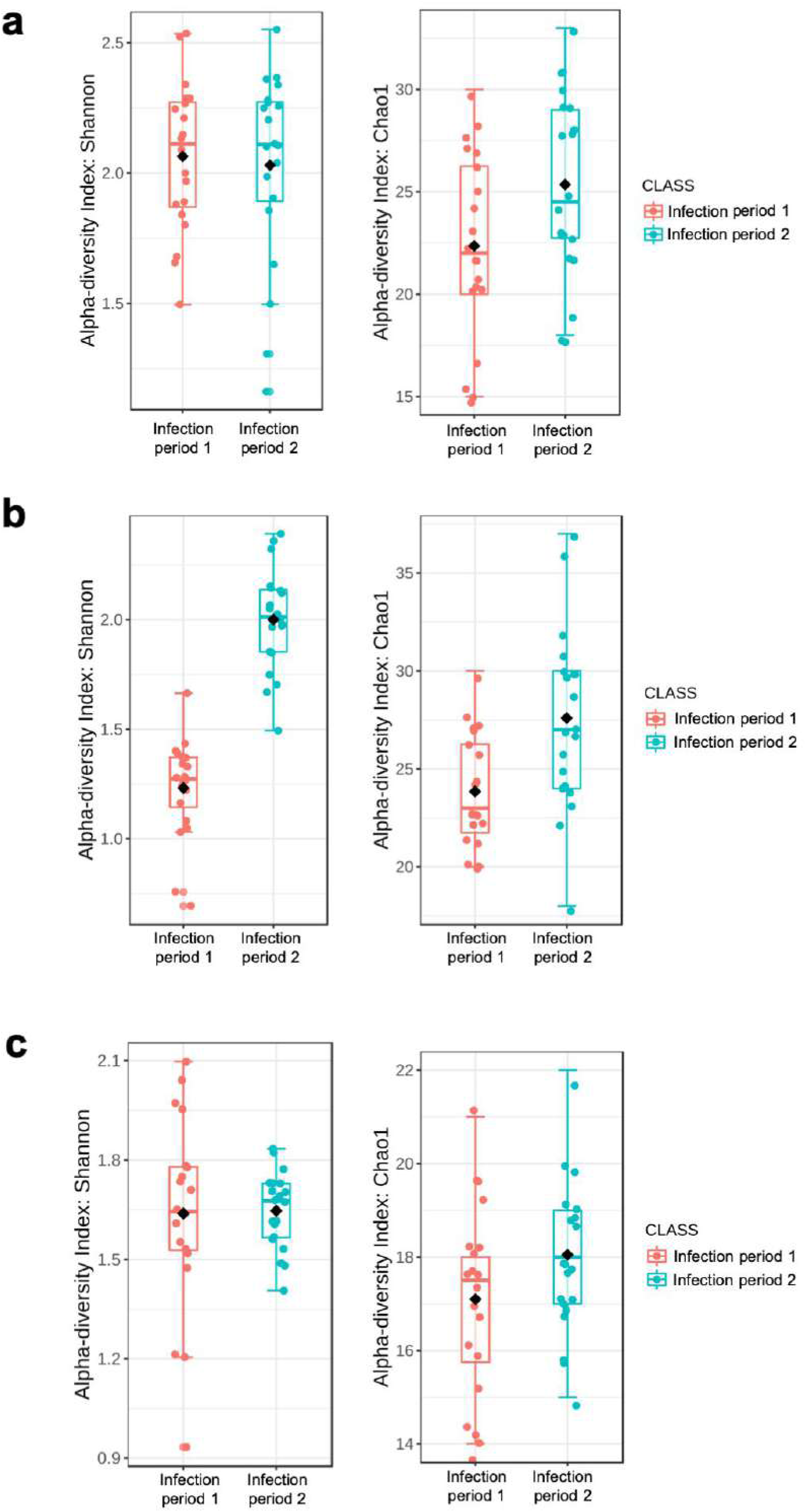
Boxplot illustrating the differences in Chao1 and Shannon diversity measures of the fungal communities between both infection periods in D.O. Ribeiro **(a)**, D.O. Rías Baixas **(b)**, and Valdeorras **(c)**.

The relative abundance of fungal families detected across sampling times is shown in Fig. 5. In D.O. Ribeiro, the most abundant families were Cladosporiaceae (34.2%), Dothioraceae (33.1%) and Sporidiobolaceae (6.3%) (initial microbiome); Cladosporiaceae (22.9%), Dothioraceae (18.4%) and Dermateaceae (14.6%) (infection period 1); and Cladosporiaceae (22.2%), Dothioraceae (15.1%) and Dermateaceae (14.9%) (infection period 2). In D.O. Rías Baixas, the most abundant families were Dothioraceae (81.3%), Cladosporiaceae (9.9%) and Pleosporaceae (4.4%) (initial microbiome); Dothioraceae (66.5%), Cladosporiaceae (9.4%) and Pleosporaceae (7.1%) (infection period 1); and Dothioraceae (26.9%), Pleosporaceae (20.4%) and Dermateaceae (8.3%) (infection period 2). In D.O. Valdeorras, the most abundant families were Dothioraceae (70.9%), Cladosporiaceae (23.1%) and Filobasidiaceae (1.2%) (initial microbiome); Dothioraceae (45.2%), Cladosporiaceae (17.8%) and Tremellaceae (8.5%) (infection period 1); and Dothioraceae (40.2%), Cladosporiaceae (20.9%) and Dermateaceae (12.8%) (infection period 2).

**Figure 5.**
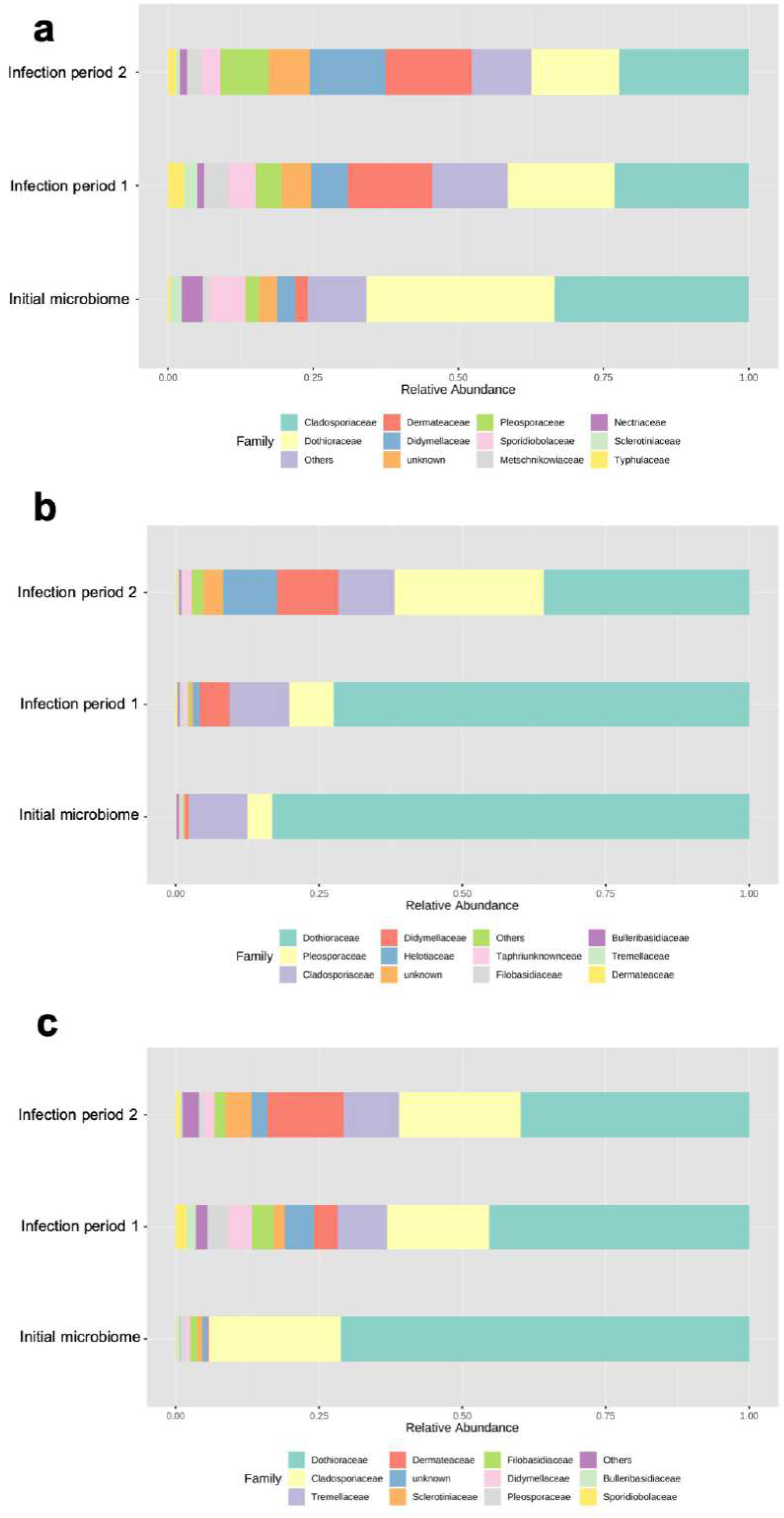
Relative abundance of different fungal families detected across sampling times (initial microbiome, infection period 1 and infection period 2) in D.O. Ribeiro **(a)**, D.O. Rías Baixas **(b)**, and Valdeorras **(c)**.

### 3.4 Infection periods specific and shared fungal assemblages

The percentage of shared fungal OTUs among the three sampling times were similar in all D.O.: 31.5% (D.O. Ribeiro), 31.4% (D.O. Rías Baixas), and 28.7% (D.O. Valdeorras) (Fig. 6). Specific OTUs associated with each sampling time ranged from 15.8 to 21.4% (D.O. Ribeiro), from 6.9 to 23.8% (D.O. Rías Baixas), and from 9.7 to 17.2% (D.O. Valdeorras). Excluding the initial fungal microbiome and comparing the two infection periods, shared fungal OTUs among infection periods were also similar: 54.1% (D.O. Ribeiro), 56.3% (D.O. Rías Baixas), and 56.0% (D.O. Valdeorras). The OTUs that were unique in both infection periods for each D.O. are shown in Table S3. Genera *Eucasphaeria* and *Penicillium* were unique to the infection period November-February, while *Cryptodiaporthe* genus was unique to the infection period February-May in the three D.O.

**Figure 6.**
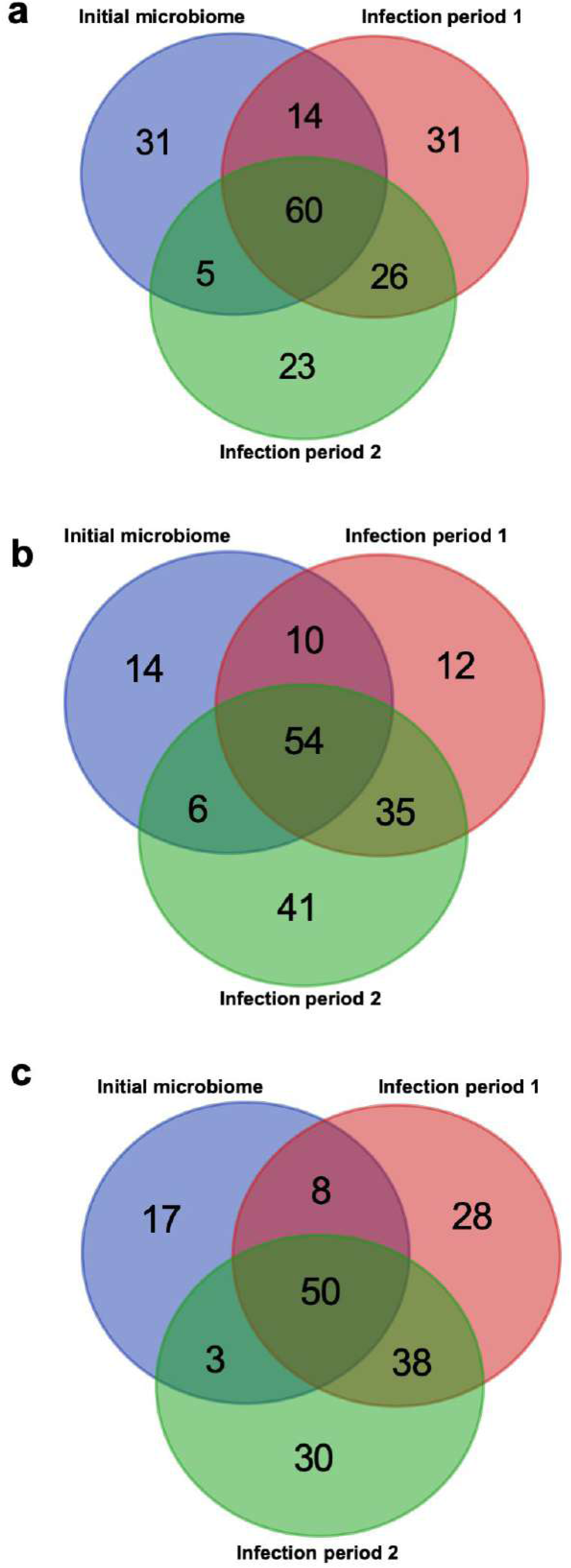
Venn diagram illustrating the overlap of the OTUs identified in the fungal microbiota among sampling times in D.O. Ribeiro **(a)**, D.O. Rías Baixas **(b)**, and Valdeorras **(c)**.

The linear discriminant analysis effect size (LEfSe) detected 3, 9 and 4 fungal clades in the grapevine inner tissues, which discriminated the fungal communities between infection periods in D.O. Ribeiro, D.O. Rías Baixas and D.O. Valdeorras, respectively (Fig. 7). The infection period 2 showed higher number of differentially abundant fungal clades (2, 8, and 3 in D.O. Ribeiro, D.O. Rías Baixas and D.O. Valdeorras, respectively). In the infection period 1, the dominant fungal genus in all D.O. was *Aureobasidium*. In the infection period 2, the dominant fungal genera were *Epicoccum* (D.O. Ribeiro), an unknown genus within the Pleospareaceae family (D.O. Rías Baixas), and *Cyanodermella* (D.O. Valdeorras).

**Figure 7.**
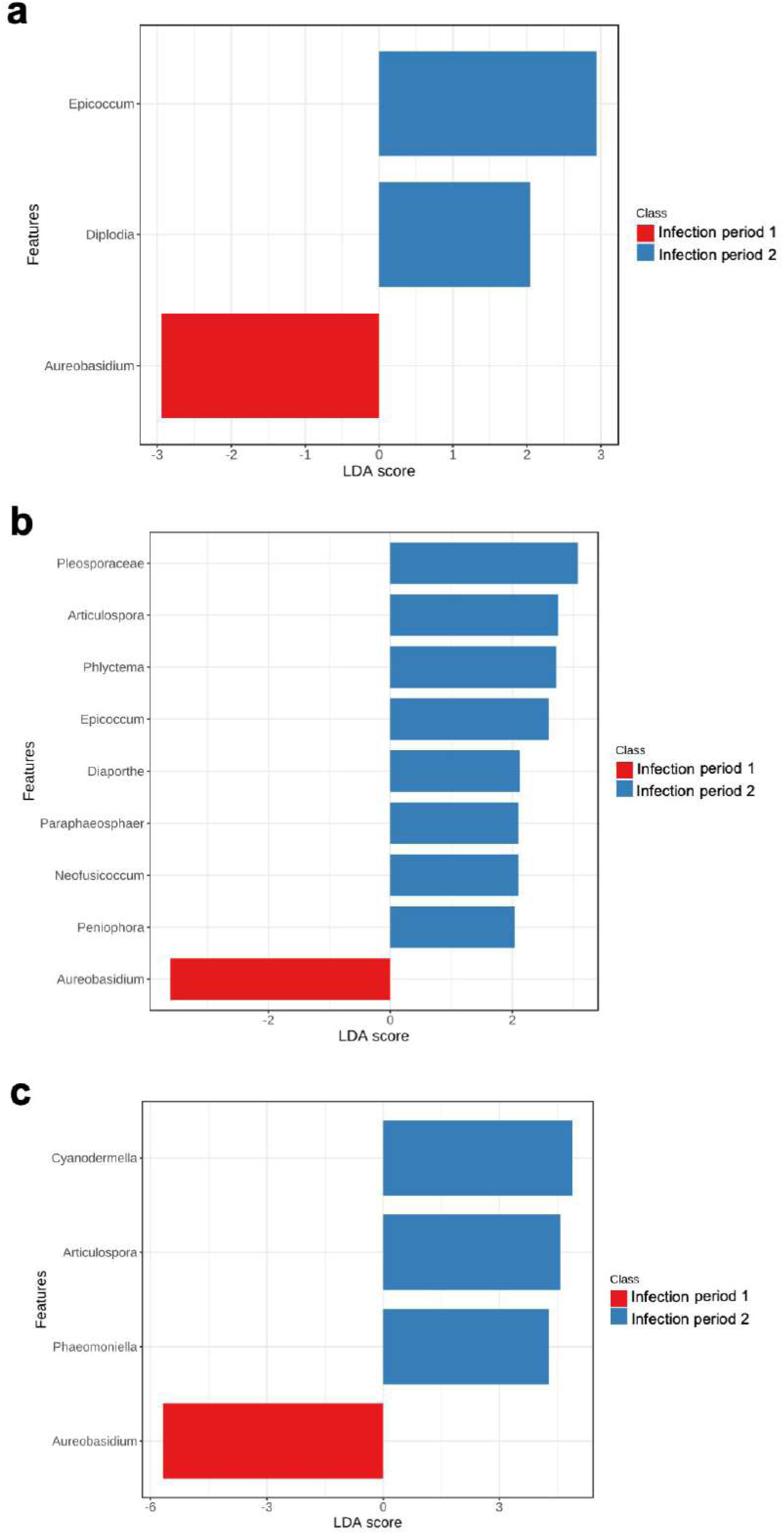
LEfSe was used to identify the most differentially abundant taxa between infection periods. Bar graph showing LDA scores for fungal genera. Only taxa meeting an LDA significant threshold >2 are shown.

### 3.5 The natural infection rates caused by fungal trunk pathogens differ between pruning times

Among the identified taxa, 10 genera are generally regarded as being associated with GTDs: *Botryosphaeria*, *Cadophora*, *Cryptovalsa*, *Cytospora*, *Diaporthe*, *Diplodia*, *Eutypa*, *Neofusicoccum*, *Phaeoacremonium* and *Phaeomoniella*. Alpha-diversity of fungal communities associated with GTDs in grapevine wood samples did not differ significantly among D.O. (Chao1: *P* = 0.1328, Shannon: *P* = 0.7608; Fig. S7). The infection periods predicted the summary metrics of alpha-diversities in D.O. Ribeiro (Chao1: *P* = 0.041, Shannon: *P* <0.001) and D.O. Rías Baixas (Chao1: *P*<0.001, Shannon: *P*<0.001), richness and diversity being higher in the infection period 2 (Fig. 8a and 8b). The alpha-diversity of fungal GTD communities did not differ between infection periods in D.O. Valdeorras (*P*>0.05; Fig. 7c).

**Figure 8.**
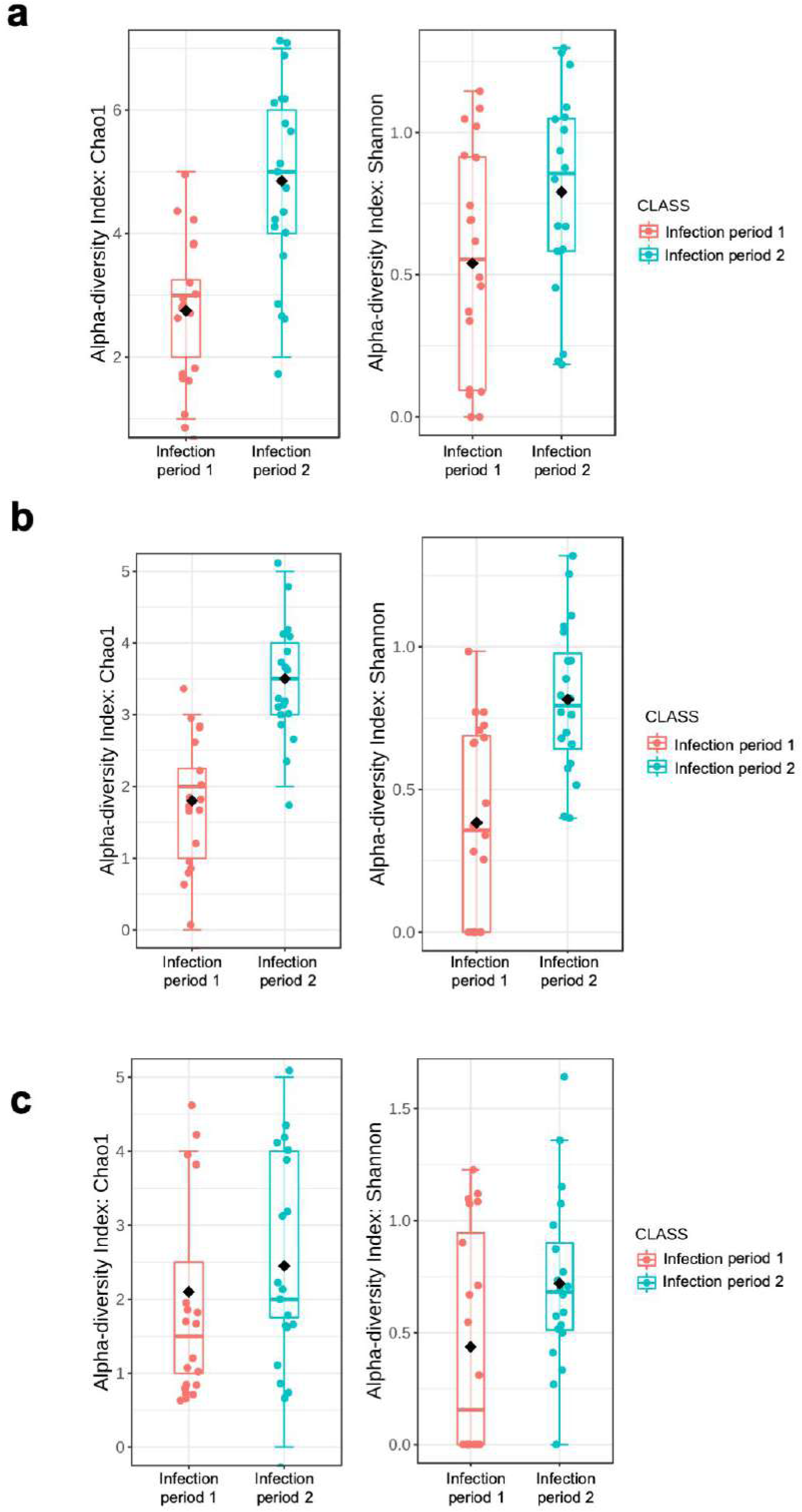
Boxplot illustrating the differences in Chao1 and Shannon diversity measures of the grapevine trunk disease pathogens between both infection periods in D.O. Ribeiro **(a)**, D.O. Rías Baixas **(b)**, and Valdeorras **(c)**.

In the annual shoot (November: initial fungal microbiome), the percentages of fungal GTD abundances with respect to the total fungal microbiome ranged from 0.1 to 0.7% (Fig. S8). Regarding the infection periods, the percentages of fungal GTD abundances with respect to the total fungal microbiome ranged from 0.2 to 1.2% (infection period 1) and from 0.3 to 1.9% (infection period 2) (Fig. 9). The abundances of several fungal GTD genera increased significantly in the infection period 2 compared to the infection period 1 (*P*<0.05; Fig. 9): *Cadophora* and *Diplodia* in D.O. Ribeiro, *Cadophora*, *Cytospora*, *Diaporthe*, *Diplodia*, *Eutypa* and *Nefosicoccum* in D.O. Rías Baixas, and *Diaporthe* and *Phaeomoniella* in D.O. Valdeorras. The abundance of *Cadophora* increased significantly in the infection period 1 compared to the infection period 2 in D.O. Valdeorras (*P*<0.05; Fig. 9c).

**Figure 9.**
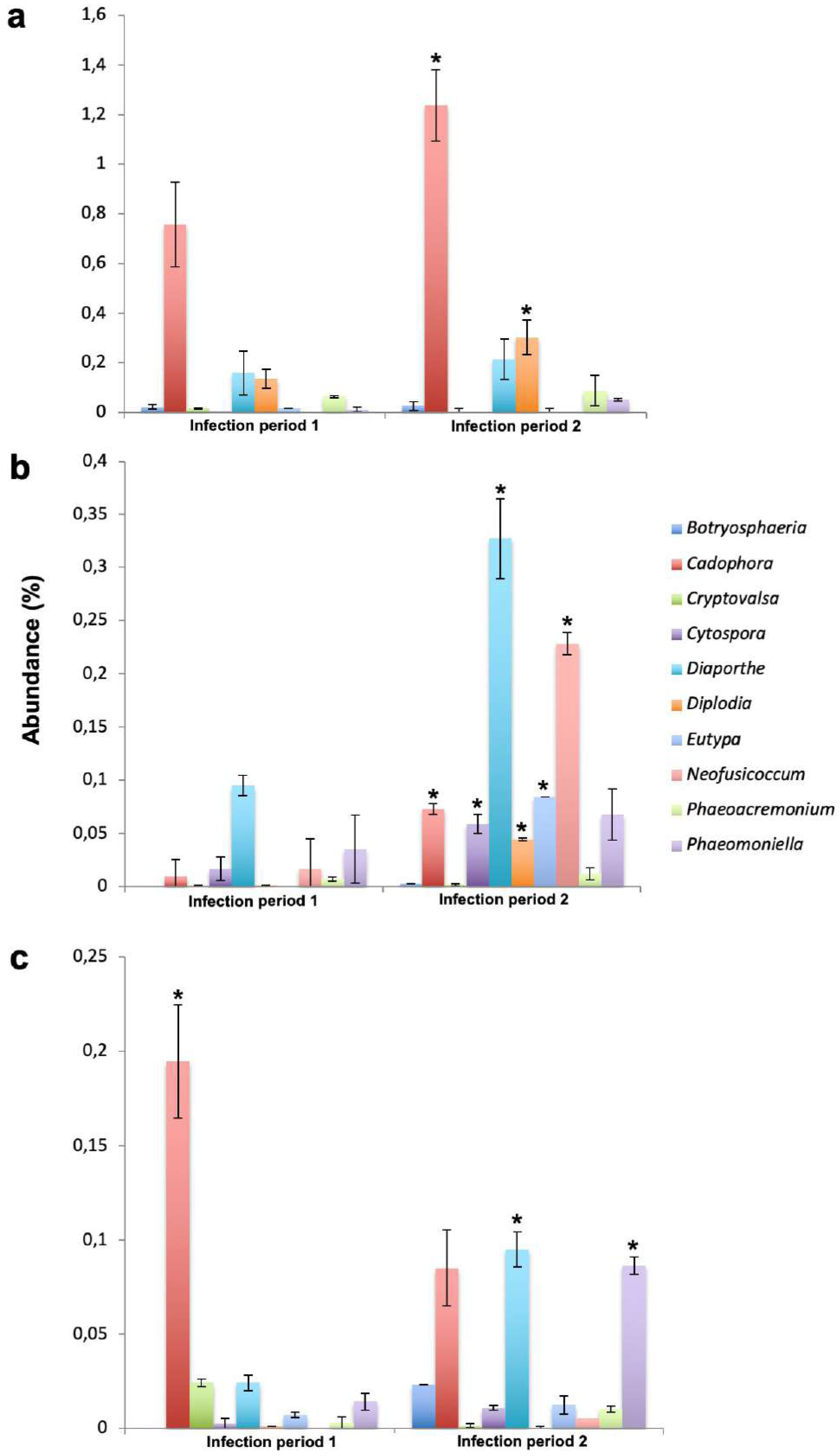
Distribution of the relative abundance of fungal trunk diseases genera obtained by high-throughput amplicon sequencing in both infection periods in D.O. Ribeiro **(a)**, D.O. Rías Baixas **(b)**, and Valdeorras **(c)**. Asterisks (*) indicate significant differences in fungal abundances between infection periods (*P*=0.05).

### 3.6 Correlation with weather variables

Climate conditions in each D.O and experimental season infection period is shown in Table S4. Climate variables varied between pruning seasons and locations. The mean values of temperature were similar during the winter season in D.O. Valdeorras (2017/2018: 6.52 ºC; 2018/2019: 7.04 ºC) and D.O. Ribeiro (2017/2018: 6.79 ºC; 2018/2019: 7.44 ºC), while they were around 3 degrees on average higher in D.O. Rías Baixas (2017/2018: 9.42 ºC; 2018/2019: 10.22 ºC). In general, temperature declined after November pruning reaching its yearly minimum during the winter season (Table S4). Temperature increased steadily from February pruning until May pruning. Accumulated rainfall was very stable after November pruning (winter season) at both D.O. Valdeorras (2017/2018: 298.40; 2018/2019: 309.60) and D.O. Ribeiro (2017/2018: 322.20 mm; 2018/2019: 294.40 mm), but it was around 100 mm on average higher in D.O. Rías Baixas (2017/2018: 393.30 mm; 2018/2019: 439.10 mm). After February pruning (spring season), this parameter increased in 2017/2018 but decreased in 2018/2019, at the three D.O. studied. In general, D.O. Rías Baixas averaged the highest rainfall among the three D.O. The relative humidity was highly stable at three locations and seasons, and as expected higher rates were recorded during winter.

A significant correlation between the main weather variables and the OTU abundances of the total fungal microbiome, *Diaporthe* and *Phaeomoniella* was detected (Table 2). Average daily temperature for the 8-week period after pruning was negatively correlated (*P*<0.05) with the OTU abundances of the total fungal microbiome. Accumulated rainfall over 8 and 11 weeks positively correlated with the fungal microbiome abundances (*P*<0.05). Regarding GTD fungal genera, a negative correlation with temperature (*P*<0.05) was observed for *Diaporthe* and *Phaeomoniella* in the first week after pruning. Accumulated rainfall over 8 and 11 weeks positively correlated with *Diaporthe* abundances (*P*<0.05) (Table 2).

**Table 2.** Spearman’s correlation coefficients of the relationships between weather data and OTUs number of the total fungal microbiome, *Cadophora, Diaporthe, Diplodia, Phaeoacremonium* and *Phaeomoniella*. All OTU data are log transformed.

| Correlation coefficient and significance | Fungal microbiome |  |  | *Cadophora* |  |  | *Diaporthe* |  |  | *Diplodia* |  |  | *Phaeoacremonium* |  |  | *Phaeomoniella* |  |  |
| --- | --- | --- | --- | --- | --- | --- | --- | --- | --- | --- | --- | --- | --- | --- | --- | --- | --- | --- |
|  | T1* | RH1 | LCR1 | T1 | RH1 | LCR1 | T1 | RH1 | LCR1 | T1 | RH1 | LCR1 | T1 | RH1 | LCR1 | T1 | RH1 | LCR1 |
| r | **-0.938** | 0.143 | 0.306 | -0.146 | 0.262 | -0.067 | **-0.599** | 0.303 | 0.161 | -0.380 | 0.303 | 0.161 | -0.068 | 0.102 | -0.298 | **-0.731** | 0.102 | -0.298 |
| P | **<0.01** | 0.658 | 0.333 | 0.649 | 0.409 | 0.833 | **0.039** | 0.337 | 0.676 | 0.229 | 0.337 | 0.676 | 0.831 | 0.750 | 0.346 | **<0.01** | 0.750 | 0.346 |
|  | **T2** | **RH2** | **LCR2** | **T2** | **RH2** | **LCR2** | **T2** | **RH2** | **LCR2** | **T2** | **RH2** | **LCR2** | **T2** | **RH2** | **LCR2** | **T2** | **RH2** | **LCR2** |
| r | **-0.754** | 0.103 | -0.188 | -0.148 | 0.251 | -0.224 | -0.372 | 0.265 | -0.017 | -0.212 | 0.254 | -0.430 | 0.066 | 0.161 | 0.328 | -0.382 | -0.074 | 0.006 |
| P | **0.004** | 0.751 | 0.558 | 0.646 | 0.414 | 0.492 | 0.231 | 0.404 | 0.955 | 0.507 | 0.424 | 0.162 | 0.838 | 0.617 | 0.282 | 0.219 | 0.818 | 0.984 |
|  | **T4** | **RH4** | **LCR4** | **T4** | **RH4** | **LCR4** | **T4** | **RH4** | **LCR4** | **T4** | **RH4** | **LCR4** | **T4** | **RH4** | **LCR4** | **T4** | **RH4** | **LCR4** |
| r | **-0.819** | -0.048 | 0.494 | -0.213 | 0.190 | 0.170 | -0.436 | 0.022 | 0.522 | -0.341 | 0.129 | -0.303 | 0.071 | 0.027 | 0.014 | -0.487 | -0.242 | 0.268 |
| P | **<0.01** | 0.882 | 0.102 | 0.505 | 0.552 | 0.596 | 0.156 | 0.944 | 0.081 | 0.277 | 0.689 | 0.338 | 0.824 | 0.933 | 0.966 | 0.107 | 0.447 | 0.398 |
|  | **T8** | **RH8** | **LCR8** | **T8** | **RH8** | **LCR8** | **T8** | **RH8** | **LCR8** | **T8** | **RH8** | **LCR8** | **T8** | **RH8** | **LCR8** | **T8** | **RH8** | **LCR8** |
| r | **-0.634** | 0.777 | **0.733** | -0.092 | 0.070 | 0.166 | -0.244 | 0.165 | **0.708** | -0.197 | 0.127 | 0.447 | 0.137 | -0.188 | -0.002 | -0.327 | -0.160 | 0.481 |
| P | **0.027** | 0.580 | **0.006** | 0.775 | 0.827 | 0.605 | 0.443 | 0.608 | **<0.01** | 0.588 | 0.692 | 0.145 | 0.670 | 0.557 | 0.847 | 0.299 | 0.619 | 0.113 |
|  | **T11** | **RH11** | **LCR11** | **T11** | **RH11** | **LCR11** | **T11** | **RH11** | **LCR11** | **T11** | **RH11** | **LCR11** | **T11** | **RH11** | **LCR11** | **T11** | **RH11** | **LCR11** |
| r | -0.280 | -0.242 | **0.685** | -0.073 | 0.070 | 0.066 | 0.005 | 0.344 | **0.822** | -0.019 | 0.018 | 0.441 | 0.481 | -0.028 | -0.071 | -0.310 | 0.473 | 0.493 |
| P | 0.337 | 0.940 | **0.013** | 0.820 | 0.828 | 0.838 | 0.986 | 0.915 | **<0.01** | 0.953 | 0.955 | 0.518 | 0.113 | 0.929 | 0.8253 | 0.326 | 0.120 | 0.103 |

T, mean daily temperature; RH, mean daily relative humidity; LCR, logarithm of accumulated rainfall.
\*Numbers following abbreviations of T, RH and LCR refer to the weather data summarized at 1, 2, 4, 8 and 11 weeks of the experimental periods in all seasons. Significant values ( $P \leq 0.05$ ) are shown in bold.

## 4. Discussion

In this study, we characterized the fungal community composition that colonizes grapevine pruning wounds at two pruning times in six vineyards belonging to three D.O. in Spain. The fungal microbiome across the three D.O. was largely composed by Ascomycota, followed by Basidiomycota. The predominant fungal phylum found in this work is consistent with the results obtained in other studies that explored the grapevine vascular tissue by culture dependent (González and Tello, 2011; Hofstetter et al., 2012; Pancher et al., 2012; Bruez et al., 2014, 2016, 2017; Dissanayake et al., 2018; Eichmeier et al., 2018; Kraus et al., 2019) or by HTAS (Dissanayake et al., 2018; Eichmeier et al., 2018; Deyett and Rolshausen, 2019, 2020; Martínez-Diz et al., 2019b) approaches. The core microbiome included the ubiquitous, fast-growing fungi *Aureobasidium* (Dothioraceae), *Cladosporium* (Cladosporiaceae), *Neofabraea* (Dermateaceae) and *Epicoccum* (Didymellaceae). This result is in line with recent studies aiming to decipher the fungal microbiome that resides in the xylem vessels of healthy grapevine branches in Germany (Kraus et al., 2019), and in the sap of grapevine under high Pierce’s disease pressure in California (Deyett and Rolshauen, 2019).

The results obtained in D.O. Rías Baixas showed a significant fraction of variation in fungal diversity (both the alpha and beta-diversity) that could be attributed to the infection period. It is interesting to note that fungal richness and diversity obtained in the infection period November-February was high relative to the period February-May in all D.O. In Mediterranean climates, drier and colder conditions usually occur after early pruning in mid-autumn, while wetter and warmer conditions favourable for fungal growth and infection occur progressively after pruning in late winter (Luque et al., 2014). The lack of significant trend in fungal microbiome abundances in both infection periods for all D.O. can be attributed to the Oceanic climate conditions in Galicia region, with temperate and rainy periods from autumn to spring, which may have favoured fungal spread and infection. In addition, two factors could also contribute to the high abundance of microbial infection during November-February, namely the wound healing and the bleeding processes. The wound healing involves the drying of the cane tissues below the pruning wounds (Bostock and Stermer, 1989), which results in a dead wood area called the drying cone (Lafon, 1921). In late winter and early spring, environmental conditions are favourable for a rapid wound healing. When the weather is cold, pruning wounds heal slowly leaving them open to fungal infection. In addition, bleeding of sap from the cut ends of canes or spurs is the first sign of renewed activity. Bleeding alone might provide some wound protection by flushing away fungal spores in early spring.

Spores are usually spread from sexual or asexual structures by wind, rain droplets or arthropods, until they land on freshly and susceptible pruning wounds and with conditions of optimal air temperature and moisture begin to germinate (Bettiga, 2013). In this study, the correlation coefficients calculated between the mean daily temperature or the accumulated rainfall and fungal microbiome infections showed negative values for temperature until eight week after pruning, and positive and statistically significant correlations for rainfall at 8 and 11 weeks after pruning. An explanatory hypothesis for the negative correlations with temperature variable might be related with a combination of favourable climatic conditions promoting a faster and suitable pruning wound healing, which physically impeded the entrance of fungal spores into the grapevine vascular tissue. Pruning grapevines in dry and warm weather is known to enhance the mechanisms which reduce pruning wounds susceptibility (Munkvold and Marois, 1995; Rolshausen et al., 2010). However, further research is required to confirm this hypothesis. Positive correlations with accumulated rainfall could indicate that rain events have an effect in increasing fungal microbiome abundance, and hence, pruning wounds infections. Several studies found that spore release and airborne inoculum spread of fungal trunk pathogens in vineyards coincided with the beginning and/or after periods of rain or irrigation events (Pearson, 1980; Carter, 1991; Michailides and Morgan, 1993; Eskalen and Gubler, 2001; Gubler et al., 2005; Amponsah et al., 2009; Kuntzmann et al., 2009; Trouillas and Gubler, 2009; Úrbez-Torres et al., 2010a; van Niekerk et al., 2010; Baskarathevan et al., 2013; Gubler et al., 2013; Úrbez-Torres et al., 2019). It has also been reported that rain can likely contribute to pycnidia and conidia masses development (Anco et al., 2013; Onesti et al., 2017), and to the splash-dispersal of conidia from pycnidia (González-Domínguez et al., 2020).

The linear discriminant analysis effect size detected several fungal clades, which discriminated the fungal communities between infection periods. The fungal genus *Aureobasidium* was predominant during the period November-February. Species of this genus, in particular *A. pullulans*, is known to dominate the microbial consortia of grapevine (Sabate et al., 2002; Martini et al., 2009; González and Tello, 2011; Barata et al., 2012; Pinto et al., 2014; Dissanayake et al., 2018; Deyett and Rolshausen, 2019; Martínez-Diz et al., 2019b). *A. pullulans* has evidenced great capacity to colonize grapevine pruning wounds (Munkvold and Marois, 1993) and to act as a biocontrol agent of several grapevine post-harvest diseases (Schena et al., 2002; Martini et al., 2009). This yeast-like fungus also showed antagonistic abilities against *Eutypa lata*, the main causal agent of Eutypa dieback of grapevine, reducing of up to 50% fungal infection in pruning wounds (Munkvold and Marois, 1993). In a recent study, *A. pullulans* reduced the *in vitro* mycelial growth of *Diplodia seriata*, one of the causal agents of Botryosphaeria dieback of grapevine, but no significant reduction of necrotic lesions were found in grapevine cuttings (Pinto et al., 2018).

Several fungal genera associated with GTDs, such as *Cadophora*, *Cytospora*, *Diaporthe*, *Diplodia* and *Phaeomoniella*, were mostly identified during the infection period February-May and explained the differences observed between periods. Cross-infection throughout both periods was unlikely to occur given the long wood section of approximately 15-cm left between sampling periods. Using artificial inoculations with extreme disease pressure, the farthest downward growth for a fast-growing fungus such as *E. lata* was estimated to be 4 cm at 5 months after inoculation (Weber et al., 2007), and the overall mean of the GTD pathogens *D. seriata* and *Phaeomoniella chlamydospora* recovery five months after inoculation were 54.2% and 46.9%, respectively, at 4.5 cm below the pruning wound (Elena and Luque, 2016). Noticeably, low GTD fungal abundance were detected in annual shoots. The data support the evidence that these fungi prefer perennial woody stems, which is where wood symptoms associated with GTDs are commonly found (Gramaje et al., 2018).

Trunk disease fungi are mainly spread through aerially dispersed spores infecting grapevines via pruning and/or natural wounds (Rolshausen et al., 2010; van Niekerk et al., 2011; Gramaje et al., 2018). Spore release varies throughout the growing season depending on the fungal pathogen, geographical location and environmental conditions (Larignon and Dubos, 2000; Eskalen and Gubler, 2001; Quaglia et al., 2009; Úrbez-Torres et al. 2010a, 2010b; van Niekerk et al., 2010; Billones-Baaijens et al., 2018; González-Domínguez et al., 2020), so information related to the dispersal patterns of GTD pathogens are indispensable to identify high-risk infection periods and to guide growers in timing management practices such as pruning time. In this sense, González-Dominguez et al. (2020) recently developed a model to predict disease risk caused by *Pa. chlamydospora* in vineyards and estimated that the pathogen dynamics were best explained when time was expressed as hydro-thermal time accounting for the effects of both temperature and rain. In the present study, evolution patterns of the correlation coefficients between weather data and OTUs abundance of GTD pathogens have been irregular with negatively and positively values being rarely statistically significant. In the first week after pruning, temperature was negatively correlated with *Diaporthe* and *Phaeomoniella* genus abundances and as previously discussed for the fungal microbiome, this fact could be associated with a mixture of proper climatic conditions favouring the pruning wound healing process. Negative correlations values between mean daily temperature and *D. seriata* and *Pa. chlamydospora* natural infections were also found in the first weeks after pruning by Luque et al. (2014). Accumulated rainfall was found to have a positive significant correlation with *Diaporthe* from eight weeks highlighting again the role of rain events in the infection and development of GTDs fungal pathogens, as earlier considered for the fungal microbiome. This same trend was also observed by Luque et al. (2014) for natural infections caused by *D. seriata*, *Pa. chlamydospora* and species of Diatrypaceae in Catalonian vineyards. Susceptibility of grapevine pruning wounds to trunk pathogens have been studied through artificial fungal inoculations in several grape-growing regions such as Australia (Ayres et al., 2016), California (Moller and Kasimatis, 1978; Munkvold and Marois, 1995; Eskalen et al., 2007; Úrbez-Torres and Gubler, 2011), France (Chapuis et al., 1998; Larignon and Dubois, 2000; Lecomte and Bailey, 2011), Italy (Serra et al., 2008), Michigan (Trese et al., 1982), South Africa (van Niekerk et al., 2011) and Spain (Elena and Luque, 2016). In general, these studies showed that wound susceptibility decreased as the period between pruning and inoculation of wounds increased, and it can be extended up to four to seven weeks for most pathogens under favourable conditions. The rate of natural infections in pruned canes (i.e., those not obtained through artificial inoculations), however, has not been extensively studied to date, and they can be estimated only through the spontaneous infections of the vines included as non-inoculated controls in artificial inoculations.

Results obtained in our study on the natural infections of pruning wounds in three D.O. in Galicia showed that higher fungal GTD infection abundances occurred more frequently in spring than in winter, thus suggesting that pruning wounds could be more susceptible to pathogens overall after a late pruning in winter. Similar results were obtained by Luque et al. (2014), who observed higher isolation percentages of several GTD fungi in culture medium following late pruning (February-May) compared with that following early pruning (November-February). In contrast, mean percentage values of natural infections caused by *Eutypa lata*were about 2% after the spring-pruning (mid-May to late June) and 13% after the winter pruning (January to February) in France (Lecomte and Bailey, 2011). Studies based on artificial inoculations also recommended late pruning to reduce GTD pathogens infections (Petzoldt et al., 1981; Munkvold and Marois, 1995; Chapuis et al., 1998; Larignon and Dubos, 2000; Eskalen et al., 2007; Serra et al., 2008, Úrbez-Torres and Gubler, 2011), although the real potential risk of infections may have been biased since these trials did not consider the presence of natural pathogenic inoculum along the experimental period.

In conclusion, a broad range of fungi was able to colonize grapevine pruning wounds at both infection periods. Pruned canes harbour a core community of fungal species, which appear to be independent of the infection period. In light of the GTD colonization results and given the environmental conditions and the geographical location of this study, early pruning is recommended to reduce the infections caused by GTD fungi during the pruning season in Galicia. It is important to note that read counts in HTAS approach are considered as semiquantitative (Amend et al., 2010). This means that there is no real quantitative relationship between spore count and read count, although a significant correlation between sequencing reads and the relative abundance of DNA of GTD fungi have been recently observed in soil samples (Berlanas et al., 2019). If precise indication of aerial spore load for one specific fungal species is required, quantitative PCR would become the tool of choice. In this sense, high-throughput droplet digital PCR protocols have been recently developed for absolute quantification of GTD fungi from environmental samples (Holland et al., 2019; Maldonado-González et al., 2020; Martínez-Diz et al., 2020).

## Acknowledgments

Funding was provided by Ministerstvo Školství, Mládeže a Tělovýchovy (CZ.02.1.01./0.0/0.0/16_017/0002334).

## Notes

https://figshare.com/projects/Natural_fungal_infections_-_Galicia_Spain_/79113

### Competing Interest Statement

The authors have declared no competing interest.

