## Supplementary material for "Grapevine pruning time affects natural wound colonization by wood-invading fungi"

**Table S1**
Main characteristics of the six vineyards used in this study.

| | Vineyard-1 | Vineyard-2 | Vineyard-3 | Vineyard-4 | Vineyard-5 | Vineyard-6 |
|---|---|---|---|---|---|---|
| Coordinates | 42°26'06.8"N 7°00'19.3"W | 42°25'37.5"N 7°00'18.3"W | 42°21'44.2"N 8°06'56.5"W | 42°21'34.4"N 8°07'09.5"W | 42°31'22.1"N 8°44'33.7"W | 42°31'22.8"N 8°44'31.9"W |
| Location | O Barco de Valdeorras | O Barco de Valdeorras | Leiro | Leiro | Ribadumia | Ribadumia |
| Province | Ourense | Ourense | Ourense | Ourense | Pontevedra | Pontevedra |
| Denomination of Origin | Valdeorras | Valdeorras | Ribeiro | Ribeiro | Rías Baixas | Rías Baixas |
| Age | 37 | 29 | 13 | 25 | 13 | 19 |
| Rootstock | 110 Richter | 110 Richter | 196-17 Castel | 196-17 Castel | 196-17 Castel | 196-17 Castel |
| Cultivar | Godello | Godello | Mencía | Mencía | Albariño | Albariño |

**Table S2**
Estimates of number of reads, sample coverage and diversity indices at the genus level for fungal profiles.

| Sample ID* | Number of reads | Good's coverage (%) | Chao1 richness | Shannon diversity |
|---|---|---|---|---|
| VA1 | 30997 | 99.99 | 14 | 0.82 |
| VA2 | 37265 | 99.91 | 13 | 0.80 |
| VA3 | 32082 | 99.25 | 13 | 0.89 |
| VA4 | 55980 | 99.90 | 10 | 0.77 |
| VA5 | 57752 | 100 | 12 | 0.92 |
| VA6 | 44741 | 99.99 | 18 | 1.52 |
| VA7 | 75471 | 99.26 | 17 | 1.18 |
| VA8 | 102463 | 99.99 | 21 | 1.75 |
| VA9 | 85422 | 99.99 | 26 | 1.95 |
| VA10 | 110137 | 99.99 | 22 | 1.49 |
| VA11 | 157989 | 99.97 | 21 | 1.39 |
| VA12 | 151246 | 99.99 | 22 | 1.54 |
| VA13 | 144063 | 99.29 | 21 | 1.79 |
| VA14 | 112761 | 99.98 | 22 | 1.79 |
| VA15 | 126625 | 99.98 | 23 | 1.82 |
| VB1 | 21438 | 99.99 | 9 | 0.72 |
| VB2 | 42497 | 99.99 | 11 | 0.75 |
| VB3 | 29866 | 99.99 | 11 | 0.67 |
| VB4 | 52414 | 99.99 | 12 | 0.77 |

| | | | | |
|---|---|---|---|---|
| VB5 | 38232 | 99.37 | 11 | 0.69 |
| VB6 | 81483 | 99.99 | 17 | 1.82 |
| VB7 | 85432 | 99.42 | 17 | 1.49 |
| VB8 | 100993 | 99.99 | 17 | 1.40 |
| VB9 | 73418 | 99.99 | 15 | 1.44 |
| VB10 | 142701 | 99.99 | 19 | 1.76 |
| VB11 | 153446 | 99.88 | 18 | 1.36 |
| VB12 | 102367 | 99.98 | 15 | 1.35 |
| VB13 | 126372 | 99.99 | 13 | 1.45 |
| VB14 | 98737 | 99.97 | 16 | 1.26 |
| VB15 | 143041 | 99.96 | 15 | 1.59 |
| EA1 | 34336 | 99.97 | 16 | 1.39 |
| EA2 | 47911 | 99.91 | 16 | 1.15 |
| EA3 | 60592 | 99.99 | 14 | 1.08 |
| EA4 | 39445 | 99.98 | 13 | 0.96 |
| EA5 | 30599 | 99.95 | 12 | 1.41 |
| EA6 | 30950 | 99.98 | 13 | 1.40 |
| EA7 | 91545 | 99.43 | 13 | 1.62 |
| EA8 | 35994 | 99.55 | 13 | 1.62 |
| EA9 | 53704 | 99.90 | 13 | 1.82 |
| EA10 | 77680 | 100 | 16 | 2.03 |
| EA11 | 78617 | 99.99 | 16 | 1.80 |

| | | | | |
|---|---|---|---|---|
| EA12 | 110987 | 99.71 | 18 | 2.14 |
| EA13 | 110881 | 99.98 | 22 | 2.27 |
| EA14 | 113505 | 99.77 | 22 | 1.95 |
| EA15 | 120807 | 99.97 | 22 | 2.13 |
| EB1 | 30916 | 99.98 | 18 | 1.34 |
| EB2 | 44533 | 99.98 | 18 | 1.25 |
| EB3 | 65869 | 99.29 | 19 | 1.77 |
| EB4 | 57978 | 99.53 | 15 | 1.48 |
| EB5 | 62357 | 99.99 | 18 | 1.58 |
| EB6 | 49423 | 99.92 | 19 | 1.94 |
| EB7 | 62882 | 99.45 | 17 | 1.71 |
| EB8 | 39264 | 99.61 | 18 | 1.99 |
| EB9 | 123476 | 99.99 | 22 | 2.23 |
| EB10 | 95639 | 99.99 | 21 | 2.24 |
| EB11 | 99738 | 99.97 | 25 | 1.87 |
| EB12 | 109848 | 99.83 | 23 | 2.04 |
| EB13 | 139663 | 99.98 | 25 | 2.13 |
| EB14 | 91304 | 99.99 | 19 | 1.67 |
| EB15 | 87918 | 99.99 | 20 | 1.78 |
| RBA1 | 78947 | 99.99 | 12 | 0.45 |
| RBA2 | 107572 | 99.99 | 14 | 0.53 |
| RBA3 | 74950 | 99.98 | 10 | 0.86 |

| | | | | |
|---|---|---|---|---|
| RBA4 | 122474 | 99.98 | 15 | 0.85 |
| RBA5 | 95505 | 99.99 | 14 | 0.73 |
| RBA6 | 92223 | 100 | 17 | 0.98 |
| RBA7 | 50186 | 100 | 16 | 0.62 |
| RBA8 | 72830 | 99.93 | 17 | 1.23 |
| RBA9 | 107543 | 99.91 | 23 | 1.08 |
| RBA10 | 92547 | 99.78 | 15 | 0.63 |
| RBA11 | 91007 | 99.34 | 26 | 2.19 |
| RBA12 | 101431 | 99.72 | 22 | 1.92 |
| RBA13 | 122762 | 99.98 | 17 | 1.67 |
| RBA14 | 95197 | 99.88 | 16 | 1.75 |
| RBA15 | 76690 | 99.99 | 20 | 2.03 |
| RBB1 | 73988 | 99.99 | 13 | 0.82 |
| RBB2 | 107288 | 99.98 | 12 | 0.42 |
| RBB3 | 37573 | 99.98 | 14 | 0.58 |
| RBB4 | 32966 | 99.98 | 11 | 0.70 |
| RBB5 | 100935 | 99.98 | 13 | 0.57 |
| RBB6 | 71106 | 99.99 | 17 | 1.14 |
| RBB7 | 102172 | 99.99 | 18 | 1.20 |
| RBB8 | 72704 | 99.99 | 17 | 1.23 |
| RBB9 | 60729 | 99.99 | 14 | 1.08 |
| RBB10 | 133564 | 100 | 18 | 1.39 |

| | | | | |
|---|---|---|---|---|
| RBB11 | 74115 | 99.99 | 23 | 2.06 |
| RBB12 | 71541 | 99.98 | 22 | 1.80 |
| RBB13 | 120200 | 100 | 17 | 1.65 |
| RBB14 | 95079 | 99.90 | 17 | 1.75 |
| RBB15 | 108127 | 99.99 | 18 | 1.88 |
| VA16 | 22135 | 99.98 | 8 | 0.89 |
| VA17 | 25774 | 99.98 | 10 | 0.75 |
| VA18 | 32272 | 99.99 | 11 | 0.80 |
| VA19 | 31106 | 99.99 | 18 | 1.24 |
| VA20 | 42558 | 99.99 | 14 | 1.12 |
| VA21 | 36801 | 99.99 | 20 | 1.52 |
| VA22 | 46534 | 99.98 | 18 | 0.91 |
| VA23 | 26136 | 99.98 | 19 | 1.48 |
| VA24 | 25613 | 99.98 | 14 | 1.33 |
| VA25 | 37975 | 99.83 | 18 | 1.59 |
| VA26 | 55837 | 99.72 | 21 | 1.47 |
| VA27 | 45041 | 99.85 | 22 | 1.75 |
| VA28 | 36321 | 99.79 | 20 | 1.43 |
| VA29 | 34329 | 99.56 | 23 | 1.86 |
| VA30 | 73180 | 99.97 | 21 | 1.63 |
| VB16 | 20896 | 99.96 | 9 | 0.73 |
| VB17 | 33556 | 99.98 | 10 | 1.05 |

| | | | | |
|---|---|---|---|---|
| VB18 | 29260 | 99.98 | 8 | 0.63 |
| VB19 | 23805 | 99.98 | 5 | 0.63 |
| VB20 | 33390 | 100 | 8 | 0.93 |
| VB21 | 39692 | 99.99 | 17 | 1.39 |
| VB22 | 49296 | 99.99 | 17 | 1.07 |
| VB23 | 32440 | 99.98 | 21 | 1.63 |
| VB24 | 29700 | 99.98 | 14 | 1.29 |
| VB25 | 34674 | 99.98 | 14 | 1.20 |
| VB26 | 48616 | 99.98 | 16 | 1.36 |
| VB27 | 43326 | 99.98 | 17 | 1.52 |
| VB28 | 41102 | 99.99 | 16 | 1.37 |
| VB29 | 32632 | 99.95 | 12 | 1.31 |
| VB30 | 65962 | 99.90 | 15 | 1.44 |
| EA16 | 30996 | 99.99 | 9 | 1.23 |
| EA17 | 26365 | 99.99 | 11 | 0.81 |
| EA18 | 30523 | 99.99 | 15 | 1.53 |
| EA19 | 24200 | 99.98 | 13 | 1.22 |
| EA20 | 32822 | 99.81 | 16 | 1.44 |
| EA21 | 34981 | 99.78 | 20 | 1.58 |
| EA22 | 37116 | 100 | 18 | 1.96 |
| EA23 | 34795 | 99.99 | 18 | 1.48 |
| EA24 | 41678 | 99.98 | 17 | 1.76 |

| | | | | |
|---|---|---|---|---|
| EA25 | 42096 | 99.99 | 21 | 1.97 |
| EA26 | 42101 | 99.90 | 21 | 1.88 |
| EA27 | 43259 | 99.93 | 20 | 1.81 |
| EA28 | 48662 | 99.93 | 25 | 2.06 |
| EA29 | 46391 | 99.93 | 20 | 2.14 |
| EA30 | 35451 | 99.96 | 16 | 2.01 |
| EB16 | 25525 | 99.98 | 16 | 1.44 |
| EB17 | 38998 | 99.98 | 17 | 2.04 |
| EB18 | 38530 | 99.98 | 17 | 1.84 |
| EB19 | 22581 | 100 | 14 | 1.52 |
| EB20 | 36746 | 99.99 | 19 | 1.64 |
| EB21 | 46211 | 99.89 | 17 | 2.11 |
| EB22 | 44363 | 99.91 | 17 | 2.07 |
| EB23 | 46907 | 99.99 | 12 | 1.90 |
| EB24 | 62412 | 99.99 | 14 | 1.76 |
| EB25 | 45877 | 99.99 | 15 | 1.52 |
| EB26 | 29742 | 99.97 | 11 | 1.40 |
| EB27 | 39639 | 99.90 | 16 | 1.57 |
| EB28 | 23434 | 99.93 | 14 | 1.81 |
| EB29 | 43869 | 100 | 13 | 1.24 |
| EB30 | 45514 | 100 | 17 | 1.07 |
| RBA16 | 31241 | 99.56 | 14 | 0.56 |

| | | | | |
|---|---|---|---|---|
| RBA17 | 40820 | 99.91 | 9 | 0.32 |
| RBA18 | 21700 | 99.99 | 5 | 0.31 |
| RBA19 | 30113 | 99.66 | 9 | 0.67 |
| RBA20 | 42209 | 99.61 | 12 | 0.55 |
| RBA21 | 32981 | 99.45 | 13 | 0.88 |
| RBA22 | 36942 | 99.78 | 15 | 0.82 |
| RBA23 | 30676 | 99.61 | 14 | 0.92 |
| RBA24 | 32895 | 99.67 | 12 | 0.56 |
| RBA25 | 42133 | 99.45 | 16 | 1.01 |
| RBA26 | 23946 | 99.48 | 18 | 1.12 |
| RBA27 | 30540 | 99.40 | 19 | 1.42 |
| RBA28 | 33873 | 99.39 | 14 | 1.23 |
| RBA29 | 35837 | 99.54 | 15 | 1.37 |
| RBA30 | 45960 | 99.32 | 18 | 1.64 |
| RBB16 | 35758 | 99.81 | 10 | 0.36 |
| RBB17 | 27059 | 99.34 | 10 | 0.44 |
| RBB18 | 25019 | 99.48 | 11 | 0.62 |
| RBB19 | 28702 | 99.88 | 10 | 0.40 |
| RBB20 | 42310 | 99.90 | 13 | 0.57 |
| RBB21 | 38842 | 99.28 | 15 | 1.02 |
| RBB22 | 40254 | 99.32 | 14 | 0.78 |
| RBB23 | 36802 | 99.56 | 14 | 0.84 |

| | | | | |
|---|---|---|---|---|
| RBB24 | 38137 | 99.98 | 16 | 1.03 |
| RBB25 | 33501 | 99.98 | 16 | 0.77 |
| RBB26 | 34645 | 100 | 21 | 2.06 |
| RBB27 | 21287 | 99.99 | 14 | 1.77 |
| RBB28 | 32235 | 99.81 | 12 | 1.26 |
| RBB29 | 35528 | 99.94 | 16 | 1.47 |
| RBB30 | 27240 | 99.98 | 16 | 1.25 |

*VA: D.O. Valdeorras; EA: D.O. Ribeiro; RB: D.O. Rias Baixas.

**Table S3**
OTUs that were unique in both infection periods for each D.O.

| D.O. Ribeiro | | D.O. Rias Baixas | | D.O. Valdeorras | |
|---|---|---|---|---|---|
| Nov-Feb | Feb-May | Nov-Feb | Feb-May | Nov-Feb | Feb-May |
| Unknown | *Arthrinium* | *Aleurobotrys* | *Bannozyma* | *Acremonium* | *Apiognomonia* |
| *Pleosporaceae* | *Aspergillus* | *Bensingtonia* | *Botryosphaeria* | *Apiognomonia* | *Bloxamia* |
| *Athelia* | Unknown *Heliotaceae* | *Boeremia* | *Bulleromyces* | *Apiotrichum* | *Botryosphaeria* |
| *Bensingtonia* | *Ceratobasidium* | *Camptophora* | *Coniosporium* | *Athelia* | Unknown *Heliotales* |
| *Colletotrichum* | *Coniosporium* | *Clavaria* | *Coniothyrium* | *Boeremia* | *Candida* |
| *Conlarium* | *Cryptodiaporthe* | *Colacogloea* | *Constantinomyces* | *Buckleyzyma* | *Coniozyma* |
| *Craterellus* | *Cyclothyrium* | *Eucasphaeria* | *Cryptodiaporthe* | *Ceratobasidium* | *Constantinomyces* |
| *Cyanodermella* | *Dendrothyrium* | *Glomus* | *Cryptosporiopsis* | *Curvularia* | *Cryptodiaporthe* |
| *Devriesia* | *Devriesia* | *Krasilnikovozyma* | *Cyanodermella* | *Eucasphaeria* | *Cryptovalsa* |
| *Dioszegia* | Unknown *Gnomoniaceae* | *Malassezia* | Unknown *Orbiliaceae* | *Exobasidium* | *Dendrophoma* |
| *Derxomyces* | *Fellozyma* | *Metschnikowia* | Unknown *Gnomoniaceae* | *Exophiala* | *Devriesia* |
| *Erythrobasidium* | *Unknown Tremellales* | *Penicillium* | *Derxomyces* | *Flagelloscypha* | *Unknown* |
| *Eucasphaeria* | *Keissleriella* | *Phaeotremella* | *Eutypa* | *Heterocephalacria* | *Gnomoniaceae* |
| *Exobasidium* | *Kwoniella* | *Phialophora* | *Exophiala* | *Krasilnikovozyma* | Unknown *Dothideomycetes* |
| *Flagelloscypha* | *Lachnella* | *Piskurozyma* | *Flagelloscypha* | Unknown *Nectriaceae* | *Endoconidioma* |
| *Kondoa* | Unknown *Pleosporaceae* | *Reddellomyces* | Unknown *Physalacriaceae* | *Microdochium* | *Unknown* |
| *Lachancea* | *Lopthiotrema* | *Sampaiozyma* | *Fonsecazyma* | *Microstroma* | *Physalacriaceae* |
| *Lecanicilium* | *Mulderomyces* | *Septoriella* | *Herpotrichia* | *Murilentithecium* | *Gnomoniopsis* |
| Unknown *Nectriaceae* | *Neofusicoccum* | *Sporidiobolales* | *Hyalotiella* | *Neophaeocryptopus* | *Hyalotiella* |
| *Neocucurbitaria* | *Papiliotrema* | *Strigula* | *Hypocreales* | *Niesslia* | *Italica* |
| *Neophaeocryptopus* | *Plagiostoma* | | *Hypsotheca* | *Penicillium* | *Kalmusia* |
| *Penicillium* | *Raffaelea* | | *Lachancea* | *Periconia* | *Lanzia* |
| *Phaeococcomyces* | Unknown | | *Lanzia* | *Phlyctema* | *Unknown* |
| *Phialophora* | *Phanerochaetaceae* | | *Lasionectria* | *Piskurozyma* | *Cystobasidiomycetes* |
| *Pleospora* | | | Unknown *Pleosporaceae* | *Pithomyces* | *Neofusicoccum* |
| *Pseudohyphozyma* | | | *Leucosporidium* | *Praetumpfia* | *Neosetophoma* |
| *Sarocladium* | | | *Lewia* | *Rhizopus* | *Unknown* |
| *Seimatosporium* | | | *Unknown Niaceae* | *Septoriella* | *Pleosporales* |
| *Stagonospora* | | | *Microdochium* | *Sporobolomyces* | *Papiliotrema* |

| | | | | | |
|---|---|---|---|---|---|
| *Sterigmatomyces*<br>*Tygervalleyomyces*<br>*Xenoramularia* | | | *Mucor*<br>*unknownganishia*<br>*Neoacrodontiella*<br>*Neocucurbitaria*<br>*Niesslia*<br>*Periconia*<br>*Phaeococcomyces*<br>*Powellomyces*<br>*Pyrenophora*<br>*Rachicladosporium*<br>*Rhizopus*<br>*Sclerostagnospora*<br>*Seiridium*<br>*Setophaeosphaeria*<br>*Sydowia*<br>*Syncephalis*<br>*Xylopsora* | *Sugiyamaella*<br>*Tetracladium*<br>*Tremellomycetes*<br>*Triposporium*<br>*Vestigium*<br>*Xenoramularia* | *Pleospora*<br>*Reddellomyces*<br>*Rhexocercosporidium*<br>*Rhizoscyphus*<br>*Sphaceloma*<br>*Sterigmatomyces*<br>*Syncephalis* |

**Table S4**

Mean values of temperature and relative humidity, and accumulated rainfall values at 1, 2 and 3 periods in each experimental season (winter: infection period Nov-Feb or spring: infection period Feb-May) for the two years of assay (2017/2018 and 2018/2019), in the three locations studied: (A) D.O. Ribeiro (Ourense), (B) D.O. Rías Baixas (Pontevedra), and (C) D.O. Valdeorras (Ourense).

**A** **D.O. RIBEIRO**

| Season | Period | Days | Mean temperature (ºC) | Mean relative humidity (%) | Accumulated rainfall (mm) |
|---|---|---|---|---|---|
| Winter 2017/18 | 1 | 30 | 5.90 | 89.93 | 171.60 |
| | 2 | 30 | 7.77 | 89.80 | 125.60 |
| | 3 | 18 | 6.64 | 86.67 | 25.00 |
| Winter 2017/18 totals | | 78 | 6.79 | 89.13 | 322.20 |
| Winter 2018/19 | 1 | 30 | 10.15 | 89.03 | 107.40 |
| | 2 | 30 | 5.51 | 90.23 | 81.60 |
| | 3 | 40 | 6.87 | 86.20 | 105.40 |
| Winter 2018/19 totals | | 104 | 7.44 | 88.26 | 294.40 |
| **Winter totals (all years)** | | | **7.12** | **88.69** | **308.30** |
| Spring 2018 | 1 | 30 | 6.96 | 83.23 | 180.60 |
| | 2 | 30 | 9.45 | 81.43 | 284.00 |
| | 3 | 39 | 13.95 | 71.08 | 62.60 |
| Spring 2018 totals | | 99 | 10.47 | 77.90 | 527.20 |
| Spring 2019 | 1 | 30 | 10.24 | 76.60 | 88.80 |
| | 2 | 30 | 11.80 | 73.47 | 119.60 |
| | 3 | 31 | 14.75 | 70.10 | 59.20 |
| Spring 2019 totals | | 91 | 12.29 | 73.35 | 267.60 |
| **Spring totals (all years)** | | | **11.38** | **75.63** | **397.40** |

**B** **D.O. RÍAS BAIXAS**

| Season | Period | Days | Mean temperature (ºC) | Mean relative humidity (%) | Accumulated rainfall (mm) |
|---|---|---|---|---|---|
| Winter 2017/18 | 1 | 30 | 8.66 | 84.40 | 173.60 |
| | 2 | 30 | 10.57 | 88.93 | 206.20 |
| | 3 | 10 | 8.25 | 81.90 | 13.50 |
| Winter 2017/18 totals | | 70 | 9.42 | 85.99 | 393.30 |
| Winter 2018/19 | 1 | 30 | 11.60 | 88.00 | 243.40 |
| | 2 | 30 | 9.38 | 79.67 | 34.40 |
| | 3 | 35 | 9.74 | 83.03 | 161.30 |
| Winter 2018/19 totals | | 95 | 10.22 | 83.54 | 439.10 |
| **Winter totals (all years)** | | | **9.82** | **84.76** | **416.20** |
| Spring 2018 | 1 | 30 | 8.85 | 77.97 | 200.20 |
| | 2 | 30 | 10.38 | 80.17 | 218.10 |
| | 3 | 39 | 13.20 | 77.44 | 143.20 |
| Spring 2018 totals | | 99 | 11.03 | 78.42 | 561.50 |

| | | | | | |
|---|---|---|---|---|---|
| Spring 2019 | 1 | 30 | 11.71 | 76.37 | 83.60 |
| | 2 | 30 | 13.14 | 70.77 | 173.70 |
| | 3 | 31 | 14.61 | 73.87 | 87.10 |
| Spring 2019 totals | | 91 | 13.17 | 73.67 | 344.40 |
| **Spring totals (all years)** | | | **12.10** | **76.05** | **452.95** |

**C** **D.O. VALDEORRAS**

| Season | Period | Days | Mean temperature (ºC) | Mean relative humidity (%) | Accumulated rainfall (mm) |
|---|---|---|---|---|---|
| Winter 2017/18 | 1 | 30 | 6.08 | 84.17 | 133.60 |
| | 2 | 30 | 7.08 | 86.93 | 149.00 |
| | 3 | 16 | 6.30 | 81.94 | 15.80 |
| Winter 2017/18 totals | | 76 | 6.52 | 84.79 | 298.40 |
| Winter 2018/19 | 1 | 30 | 9.06 | 86.73 | 81.40 |
| | 2 | 30 | 4.76 | 86.10 | 46.40 |
| | 3 | 44 | 7.20 | 80.14 | 181.80 |
| Winter 2018/19 totals | | 104 | 7.04 | 83.76 | 309.60 |
| **Winter totals (all years)** | | | **6.78** | **84.27** | **304.00** |
| Spring 2018 | 1 | 30 | 6.22 | 76.93 | 158.60 |
| | 2 | 30 | 8.66 | 76.20 | 214.40 |
| | 3 | 39 | 13.49 | 67.85 | 55.20 |
| Spring 2018 totals | | 99 | 9.83 | 73.13 | 428.20 |
| Spring 2019 | 1 | 30 | 11.13 | 63.63 | 35.60 |
| | 2 | 30 | 11.15 | 71.63 | 148.40 |
| | 3 | 24 | 15.72 | 63.17 | 21.80 |
| Spring 2019 totals | | 84 | 12.45 | 66.36 | 205.80 |
| **Spring totals (all years)** | | | **11.14** | **69.74** | **317.00** |

**Figure S1.** Rarefaction curve values for each sample in each Denomination of Origin.

**Figure S2.** Relative abundance of different fungal phyla **(a)**, orders **(b)** and families **(c)** detected across Denominations of Origin.

**Figure S3.** Boxplot illustrating the differences in Chao1 and Shannon diversity measures of the fungal communities between vineyards in D.O. Ribeiro **(a)**, D.O. Rías Baixas **(b)**, and Valdeorras **(c)**.

**Figure S4.** Boxplot illustrating the differences in Chao1 and Shannon diversity measures of the fungal communities between years in D.O. Ribeiro **(a)**, D.O. Rías Baixas **(b)**, and Valdeorras **(c)**.

**Figure S5.** Boxplot illustrating the differences in Chao1 and Shannon diversity measures of the fungal communities among sampling times in D.O. Ribeiro **(a)**, D.O. Rías Baixas **(b)**, and Valdeorras **(c)**.

**Figure S6**. Principal Coordinate Analysis (PCoA) based on Bray Curtis dissimilarity metrics showing the distance in the fungal communities among sampling times D.O. Ribeiro **(a)**, D.O. Rías Baixas **(b)**, and Valdeorras **(c)**.

**Figure S7.** Boxplot illustrating the differences in Chao1 **(a)** and Shannon **(b)** diversity measures of the grapevine trunk disease pathogens among Denominations of Origin.

**Figure S8.** Distribution of the relative abundance of fungal trunk diseases genera obtained by high-throughput amplicon sequencing in the annual shoot (sampling in November: initial microbiome) in the three Denominations of Origin.

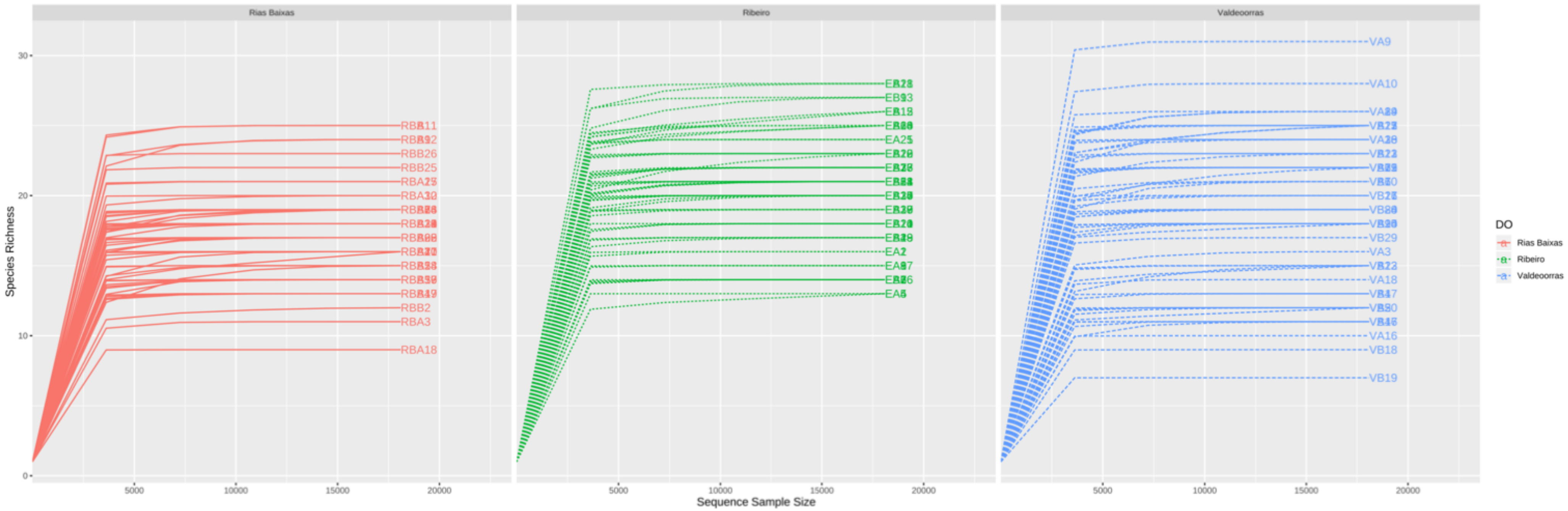

Rias Baixas
Ribeiro
Valdeoorras
Species Richness
Sequence Sample Size
DO
Rias Baixas
Ribeiro
Valdeoorras
RBB26
RBB25
RBB2
RBA3
RBA18
EA25
EA2
EA5
VA9
VA10
VB29
VA3
VA18
VA16
VB18
VB19

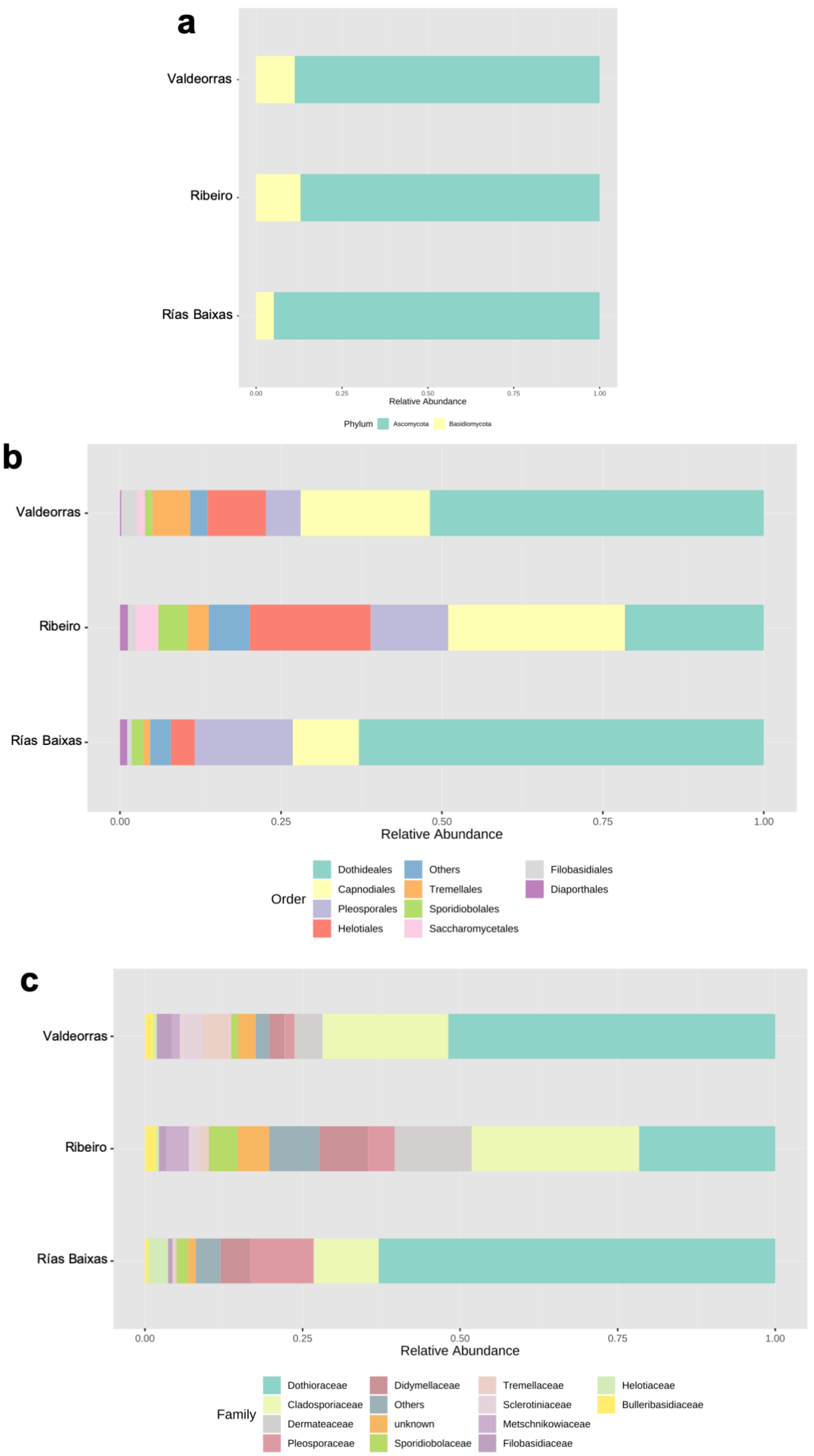

a
Valdeorras
Ribeiro
Rías Baixas
0.00
0.25
0.50
0.75
1.00
Relative Abundance
Phylum
Ascomycota
Basidiomycota
b
Valdeorras
Ribeiro
Rías Baixas
0.00
0.25
0.50
0.75
1.00
Relative Abundance
Order
Dothideales
Capnodiales
Pleosporales
Helotiales
Others
Tremellales
Sporidiobolales
Saccharomycetales
Filobasidiales
Diaporthales
c
Valdeorras
Ribeiro
Rías Baixas
0.00
0.25
0.50
0.75
1.00
Relative Abundance
Family
Dothioraceae
Cladosporiaceae
Dermateaceae
Pleosporaceae
Didymellaceae
Others
unknown
Sporidiobolaceae
Tremellaceae
Sclerotiniaceae
Metschnikowiaceae
Filobasidiaceae
Helotiaceae
Bulleribasidiaceae

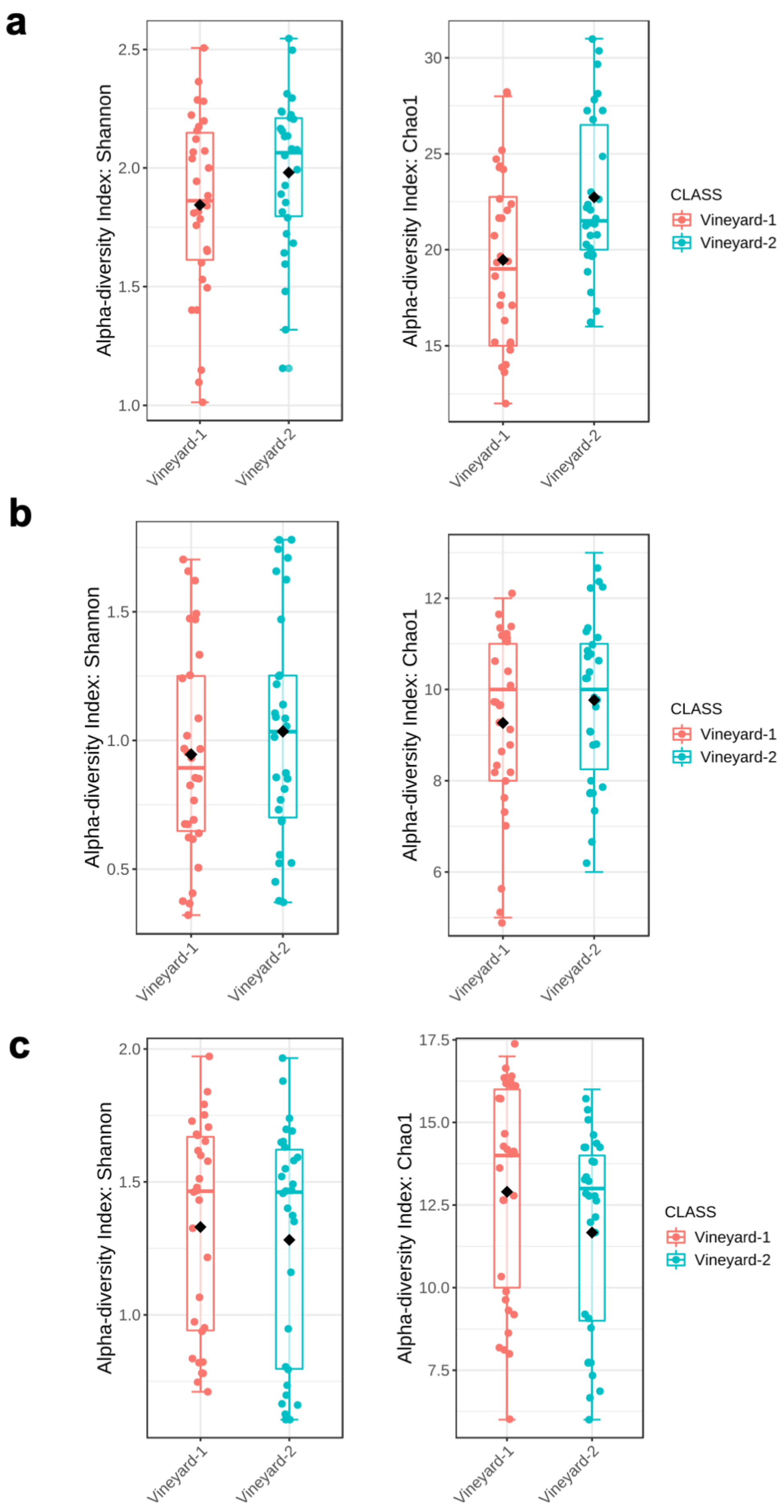
a
b
c
Alpha-diversity Index: Shannon
Alpha-diversity Index: Chao1
CLASS
Vineyard-1
Vineyard-2

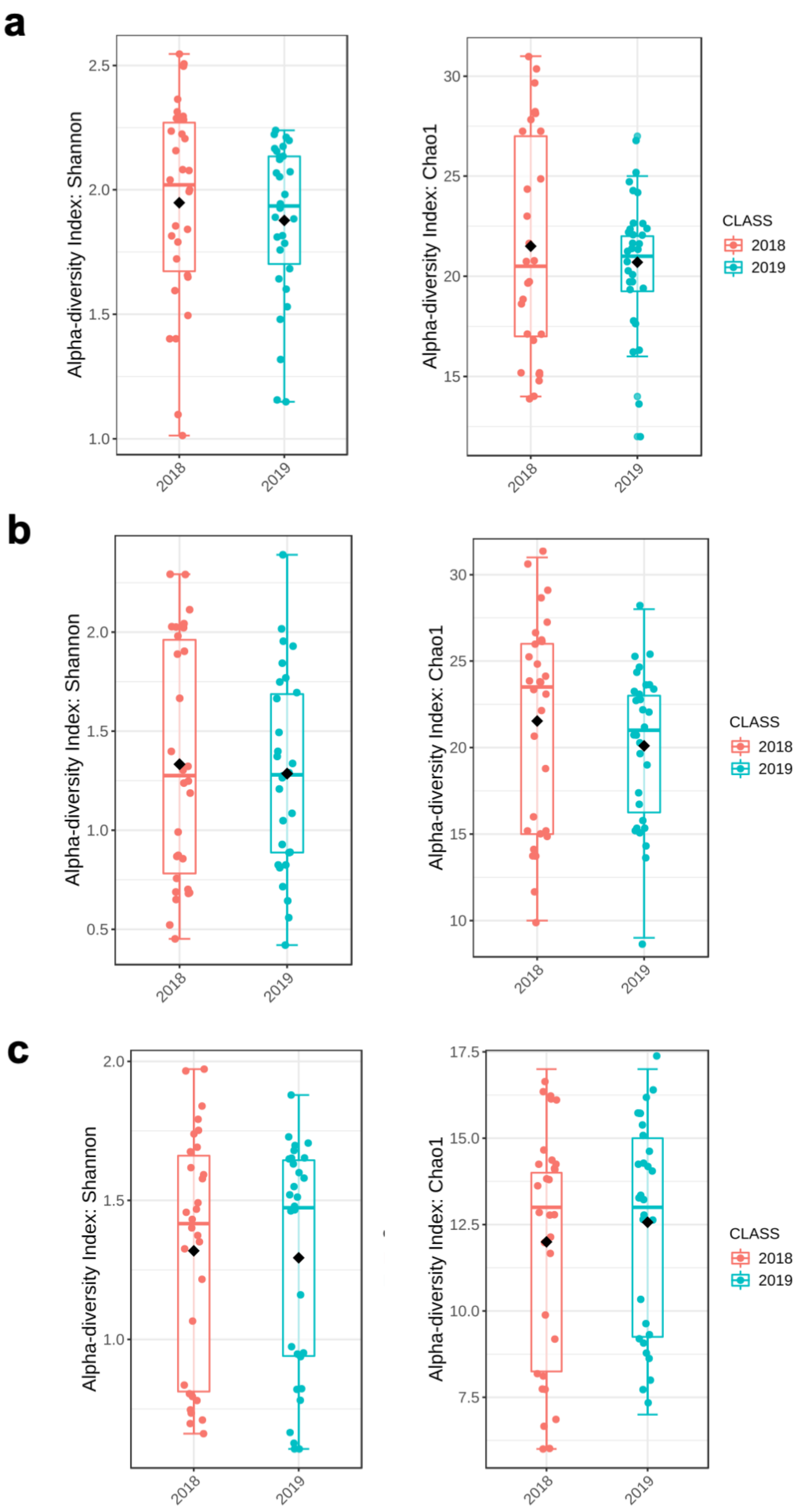

a
b
c
Alpha-diversity Index: Shannon
Alpha-diversity Index: Chao1
CLASS
2018
2019

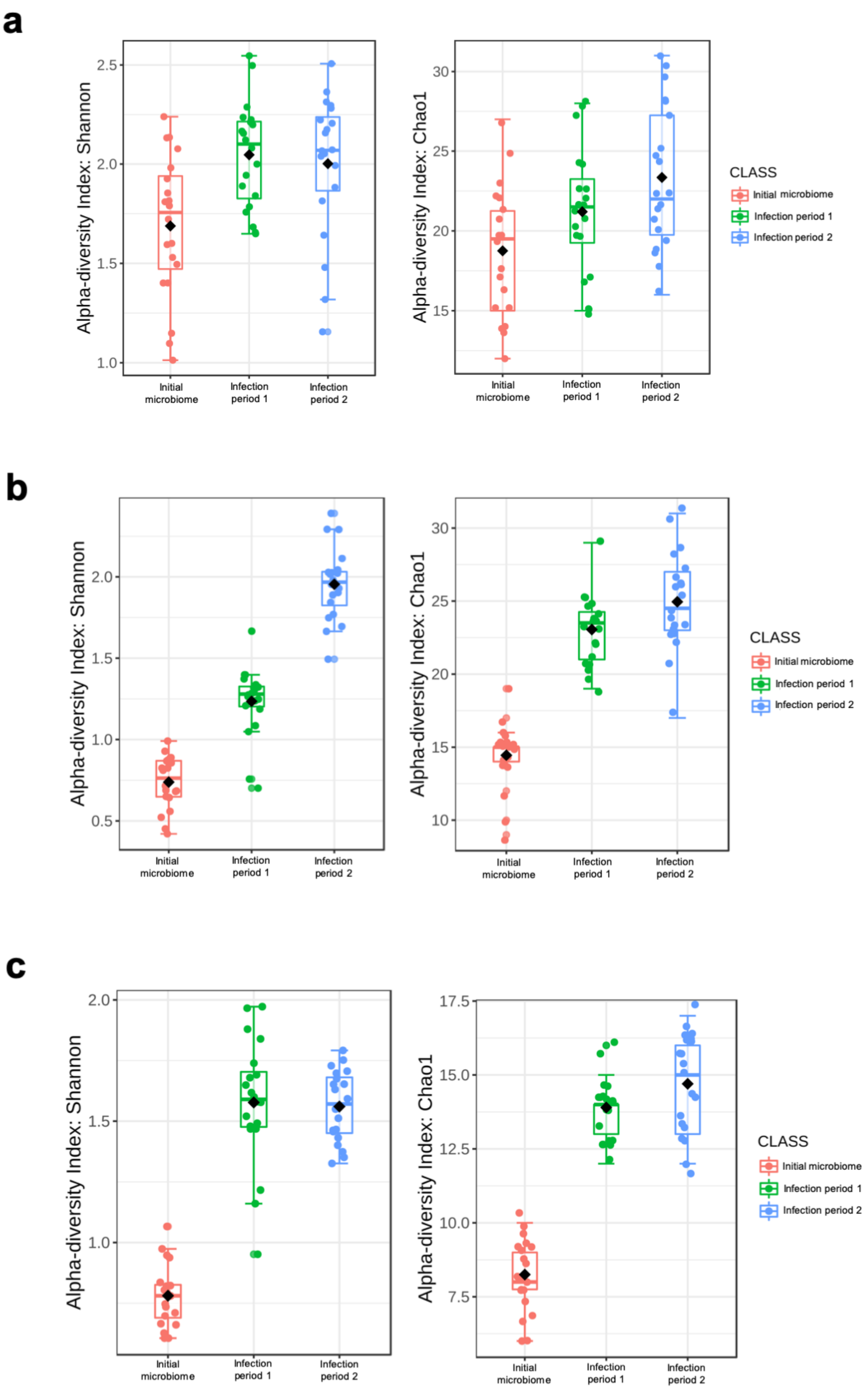
a
b
c
Alpha-diversity Index: Shannon
Alpha-diversity Index: Chao1
Initial microbiome
Infection period 1
Infection period 2
CLASS
Initial microbiome
Infection period 1
Infection period 2
1.0
1.5
2.0
2.5
0.5
15
20
25
30
10
7.5
10.0
12.5
15.0
17.5

a

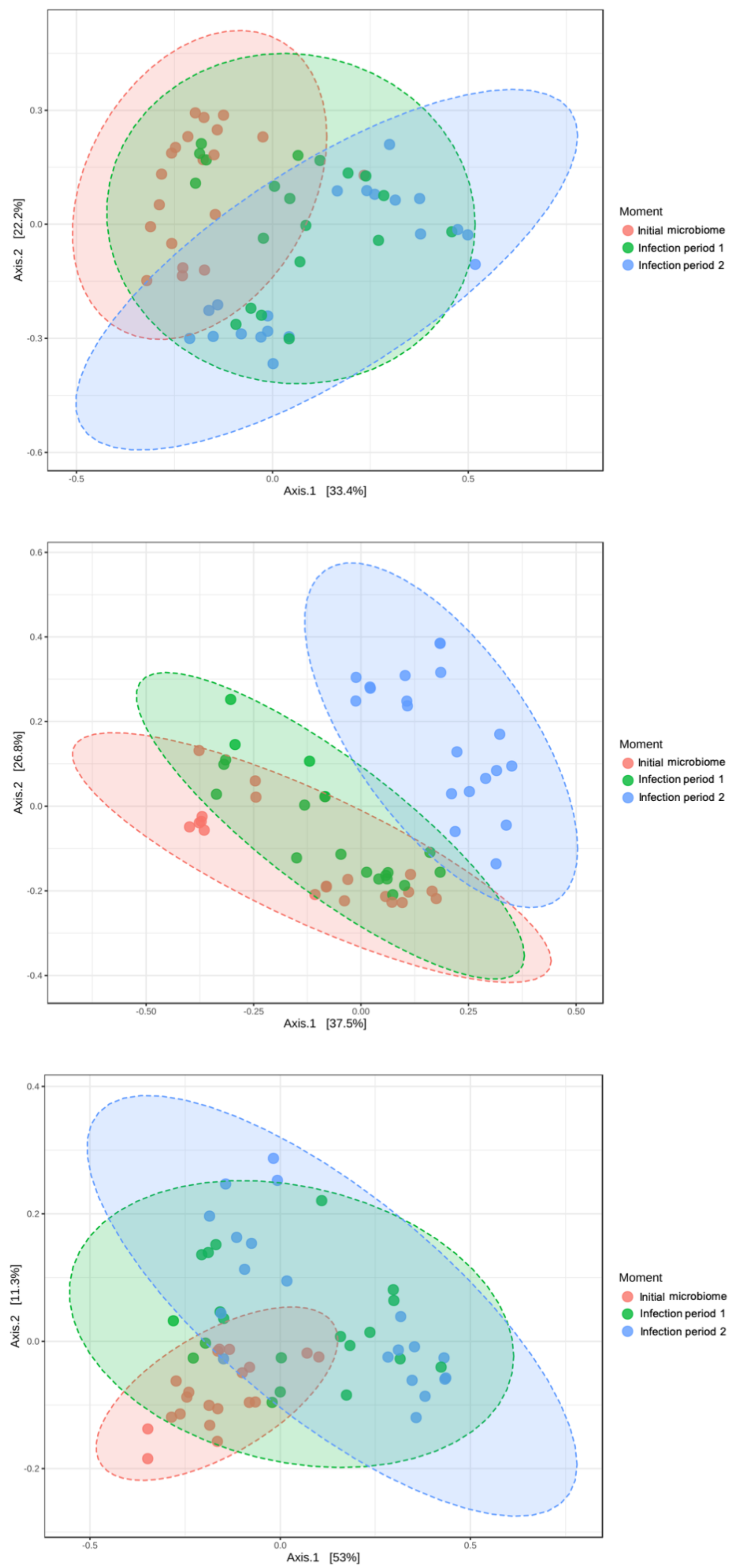
0.3
0.0
-0.3
-0.6
Axis.2 [22.2%]
-0.5
0.0
0.5
Axis.1 [33.4%]
Moment
Initial microbiome
Infection period 1
Infection period 2
b
0.6
0.4
0.2
0.0
-0.2
-0.4
Axis.2 [26.8%]
-0.50
-0.25
0.00
0.25
0.50
Axis.1 [37.5%]
Moment
Initial microbiome
Infection period 1
Infection period 2
c
0.4
0.2
0.0
-0.2
Axis.2 [11.3%]
-0.5
0.0
0.5
Axis.1 [53%]
Moment
Initial microbiome
Infection period 1
Infection period 2

**a**

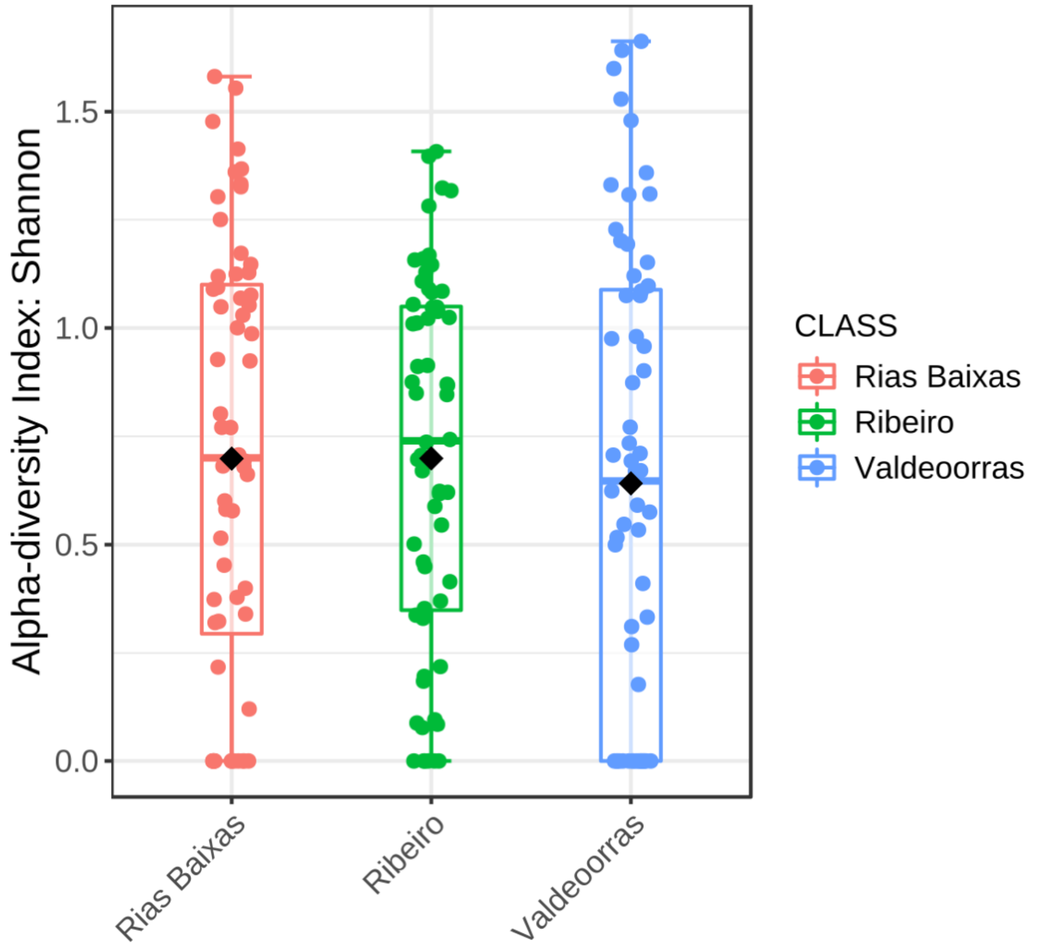

Alpha-diversity Index: Shannon
1.5
1.0
0.5
0.0
Rias Baixas
Ribeiro
Valdeoorras
CLASS
Rias Baixas
Ribeiro
Valdeoorras


**b**

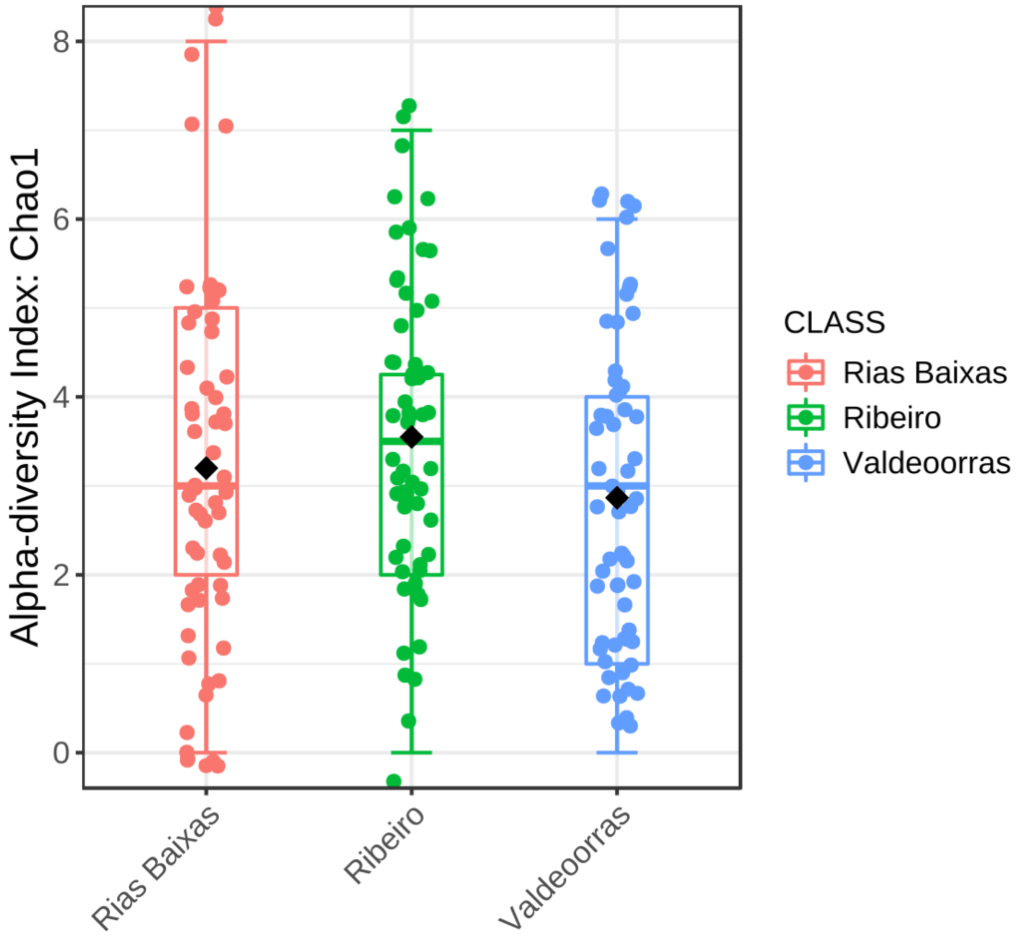

Alpha-diversity Index: Chao1
8
6
4
2
0
Rias Baixas
Ribeiro
Valdeoorras
CLASS
Rias Baixas
Ribeiro
Valdeoorras

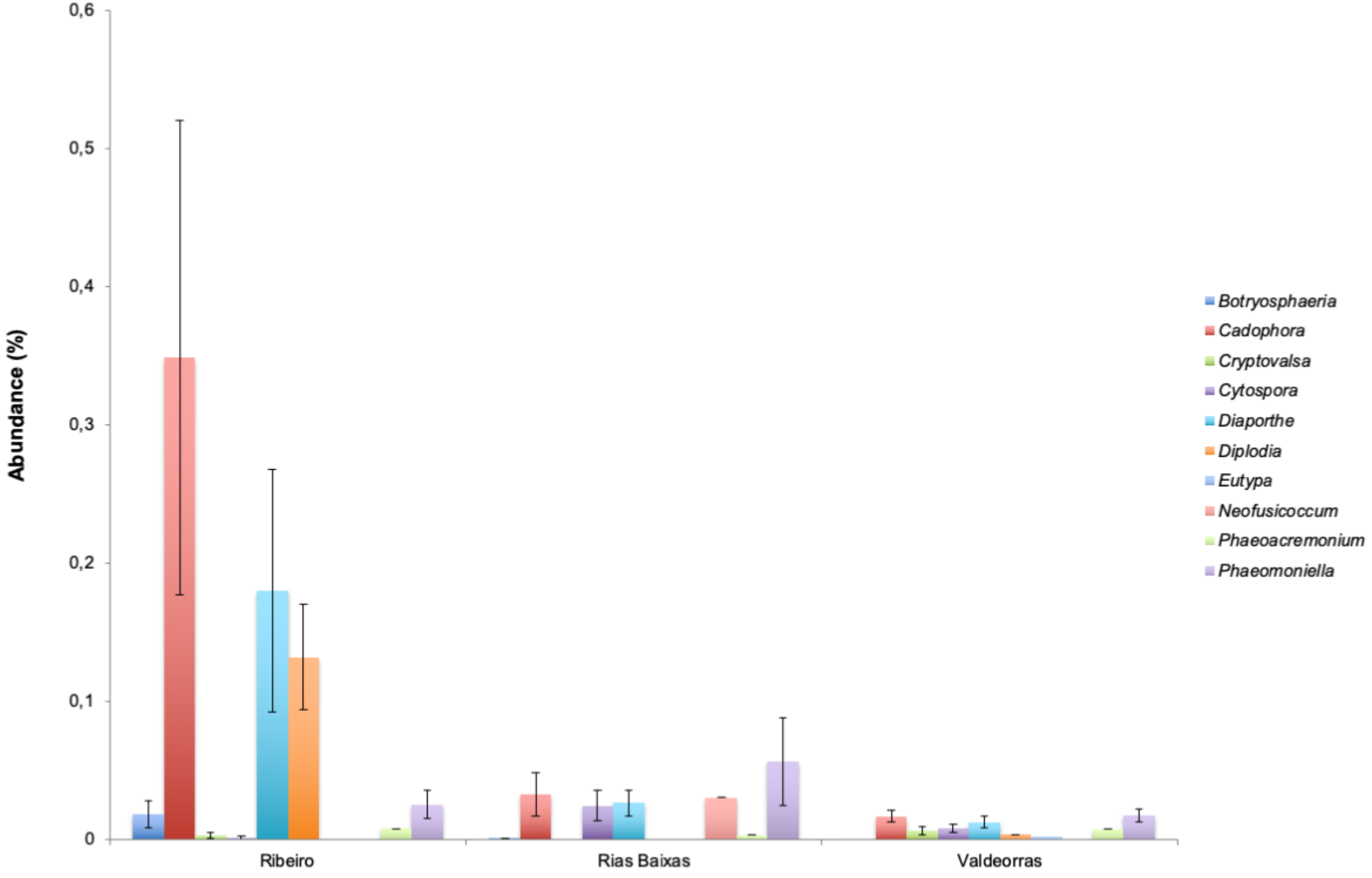
Abundance (%)
0,6
0,5
0,4
0,3
0,2
0,1
0
Ribeiro
Rias Baixas
Valdeorras
Botryosphaeria
Cadophora
Cryptovalsa
Cytospora
Diaporthe
Diplodia
Eutypa
Neofusicoccum
Phaeoacremonium
Phaeomoniella
